# The Causal Role of Beta Band Desynchronization: Individualized High-Definition Transcranial Alternating Current Stimulation Improves Bimanual Motor Control

**DOI:** 10.1101/2024.10.30.621096

**Authors:** Sybren Van Hoornweder, Diego Andres Blanco Mora, Marten Nuyts, Koen Cuypers, Stefanie Verstraelen, Raf Meesen

## Abstract

**Objective:** To unveil if 3 mA peak-to-peak high-definition β transcranial alternating current stimulation (tACS) applied over C4 –the area overlaying the right sensorimotor cortex– enhances bimanual motor control and affects movement-related β desynchronization (MRβD), thereby providing causal evidence for the polymorphic role of MRβD in motor control.

**Methods:** In this sham-controlled, crossover study, 36 participants underwent 20 minutes of fixed 20 Hz tACS; tACS individualized to peak β activity during motor planning at baseline; and sham tACS randomized over three consecutive days. Before, during, and after tACS, participants performed a bimanual tracking task (BTT) and 64-channel electroencephalography (EEG) data was measured. Spatiotemporal and temporal clustering statistics with underlying linear mixed effect models were used to test our hypotheses.

**Results:** Individualized tACS significantly improved bimanual motor control, both online and offline, and increased online MRβD during motor planning compared to fixed tACS. No offline effects of fixed and individualized tACS on MRβD were found compared to sham, although tACS effects did trend towards the hypothesized MRβD increase. Throughout the course of the study, MRβD and bimanual motor performance improved. Exclusively during motor planning, MRβD was positively associated to bimanual motor performance improvements, emphasizing the functionally polymorphic role of MRβD. tACS was well tolerated and no side-effects occurred.

**Conclusion:** Individualized β-tACS improves bimanual motor control and enhances motor planning MRβD online. These findings provide causal evidence for the importance of MRβD when planning complex motor behavior.

## 1. INTRODUCTION

Motor control is imperative to human behavior. A deeper understanding of it not only advances our fundamental knowledge of the brain but also has the potential to revolutionize therapeutic strategies for neurological conditions, such as stroke and Parkinson’s disease [1, 2]. The combination of electroencephalography (EEG) and transcranial alternating current stimulation (tACS) is promising for motor control. While EEG provides real-time monitoring of brain activity, tACS enables noninvasive modulation of neural oscillations, offering a unique ability to causally probe brain-behavior relationships [3–5].

tACS applies weak, oscillatory electrical currents to the scalp. Although incapable of generating action potentials, the resulting time-varying electric fields modulate neural activity by synchronizing endogenous rhythms to the imposed frequency, a process known as entrainment [6, 7]. According to the Arnold Tongue Hypothesis, the likelihood of successful entrainment depends on the alignment between the exogenous tACS frequency and the endogenous rhythms [8]. Besides entrainment, tACS also has neuroplasticity-like effects which can explain its after-effects [9–11]. Furthermore, when inducing low-intense electric fields, which is typically the case in humans, computational work suggests that tACS may even desynchronize ongoing neural oscillations [12]. While tACS can be applied at various frequencies, the β-band (13.5 – 30 Hz) is of particular relevance for motor behavior, as β-tACS has been shown to enhance early motor consolidation [13, 14], speed up motor performance at the cost of accuracy [15], and alter corticospinal excitability [16].

While these findings position tACS as a compelling tool to enhance motor control, little is known about its impact on event-related neural processes, which are crucial for cognition and behavior. The few studies that have explored this, indicate that tACS enhances event-related neural processes during cognitive tasks [17–19].

A key event-related feature for motor behavior is movement-related β desynchronization (MRβD), a transient β power decrease in sensorimotor regions, during the planning and execution of spontaneous, imagined and triggered movements [20–23]. The observations that motor planning MRβD is associated to force production [24], movement direction uncertainty [25] and interlimb motor control [26–28], and is attenuated in motor disorders [29, 30], has caused some to perceive MRβD as essential for motor control. However, other work indicates that MRβD might be an epiphenomenon, being insensitive to the used effector [31] and movement speed [32] during motor execution. Previously, we brought these contradictory results together by arguing that MRβD is functionally polymorphic, representing movement- and performance-specific features during planning, while representing more general unspecific processes during execution [27].

There are gaps in our understanding of tACS effects on motor control and MRβD, and the polymorphic role of MRβD in motor control. It’s also unclear whether tailoring tACS to individual β frequencies, which concurs with the Arnold Tongue hypothesis, yields neurophysiological and/or behavioral advantages over conventional fixed-frequency tACS, which is easier to implement.

Therefore, we investigated the effects of β-tACS on MRβD and the bimanual tracking task (BTT), and probed the association between MRβD and motor control. We hypothesized that tACS increases MRβD magnitude during motor planning, with individualized tACS yielding the greatest enhancement. Likewise, we expected both tACS protocols to improve bimanual motor control compared to sham, with individualized tACS being most effective. We also anticipated that motor performance improvements would relate to MRβD changes, providing further insights into motor control mechanisms and positioning tACS as a potential tool for rehabilitation.

## 2. Material and Methods

This study concurred with the Declaration of Helsinki, and was approved by the ethical committee of Hasselt University (protocol number: B1152020000017).

Thirty-six, healthy right-handed participants, aged between 18 and 30, were recruited via flyers and social media. The sample size calculations and in- and exclusion criteria are outlined in **Appendix 2**.

### 2.1. Procedure

Participants visited the lab for three consecutive days at the same hour. Each day, a different tACS condition was applied in a counterbalanced order to control for carry-over and retention effects. Due to the overt tACS artefacts in the EEG data, only participants were blinded.

Participants were then seated while their head was measured and EEG-tACS was set up. Subsequently, they were introduced to the BTT via a brief familiarization. Next, the protocol outlined in **Figure 1A** was followed: a 2-minute resting state, followed by a 5-minute baseline BTT, and 20-minutes of tACS, of which 10 minutes consisted of the BTT. After tACS, a 5-minute offline BTT block and 2-minute rest block were performed. At the end, participants were asked if they received verum or sham tACS.

**Figure 1.**
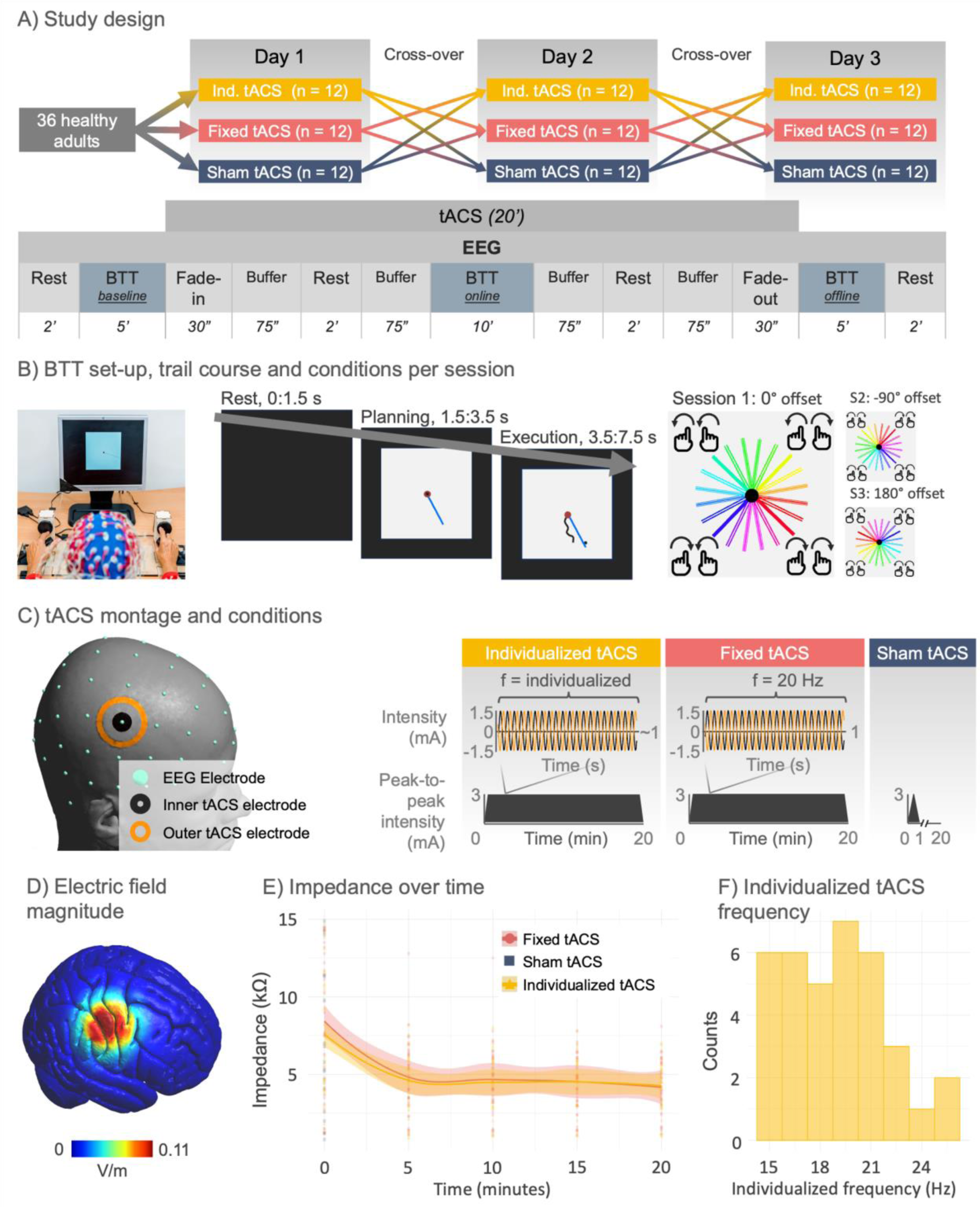
Overview of the study design and methods. **A)** Study design. Participants received 3 tACS types over 3 consecutive days. The upper part shows the condition counterbalancing, the lower part shows the content of one experimental session. Rest indicates resting-state EEG. Buffer indicates time where researchers set up the bimanual tracking task (BTT), resting-state windows and the EEG-tACS protocol. Fade-in and -out relate to the start and end of tACS. **B)** BTT: Left – Task-setup; Middle – Trial time course; Right – All BTT conditions, with 4 movement patterns (quadrants) and 5 inter-hand frequencies. Single, double- and triple lines denote both hands moving at identical speeds, or one hand moving two or three times faster. Colors denote unique conditions. In sessions 2 and 3, visual offsets were introduced to mitigate learning from the day before. **C)** tACS – EEG setup and paradigms: individualized, fixed and sham high-density tACS were applied over sensor C4. **D)** SimNIBS simulated electric field magnitude at the tACS peak intensity in the MNI head model. **E)** Impedance during fixed and individualized tACS and the initial period of sham. **F)** Histogram of individualized tACS frequencies.

### 2.2. Bimanual Tracking Task

The BTT assessed bimanual motor control (**Figure 1B**) [33]. Participants were seated in front of a screen with their forearms resting on a table, holding a fixated handlebar in each hand. Each index finger was placed in a circular groove on a rotatable dial connected to a shaft encoder registering angular displacement at 100 Hz. Participants were instructed to follow a moving target dot on the screen by simultaneously rotating the dials, with left and right rotations corresponding to cursor movements along the y- and x-axes, respectively.

Twenty unique conditions were tested, varying by inter-hand frequency and directional pattern. Five inter-hand frequencies were used; 1:1, 1:2, 2:1, 3:1 and 1:3 indicating the relative speed of the left and right hands. Four directional patterns were used: both hands moving rightward, leftward, inward or outward. Each trial consisted of three phases: rest (1.5 s), motor planning (2 s) and motor execution (4 s). During planning, participants could see the imposed line but were not allowed to move, until an auditory signal signaled the start of the execution phase.

The baseline- and offline BTT blocks consisted of 40 trials, while the online block contained 80. All 20 conditions occurred equally, with the order being randomized per block. The on-screen shown lines were visually offset by −90° and 180° rotations on days 2 and 3, respectively, to minimize retention effects across days (**Figure 1B**).

BTT performance was quantified as tracking error (TE), a measure of compliance to the imposed coordination pattern [27, 34]. TE at timepoint t was computed as:

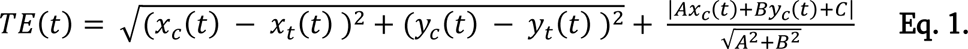

Where [*x*_*c*_(*t*), *y*_*c*_(*t*)] are the coordinates of the participants cursor at timepoint t, [*x*_*t*_(*t*), *y*_*t*_(*t*)] are the coordinates of the target dot, and *A*, *B* and *C* are the target line’s linear equation. Thus, TE informs on the participant’s cursor’s Euclidean distance to the target dot, and its orthogonal distance to the target line. The data were downsampled to 5 Hz, and TE in the online and offline block were normalized relative to baseline given our focus on TE changes following tACS:

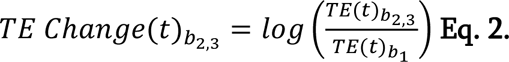

### 2.3. Transcranial Alternating Current stimulation

tACS was administered via two carbon-rubber ring electrodes connected to a neuroConn DC-Stimulator Plus. The electrodes, centered over C4, were applied using Ten20 paste (**Figure 1C and 1E**) (cf., [35]). The choice for C4 is based on previous literature indicating that β-activity in the non-dominant hemisphere is more responsive to task complexity [27, 28, 36].

tACS was delivered for 20 minutes at 3 mA (peak-to-peak) with a 30 s fade-in and fade-out period (**Figure 1D**). Fixed- and sham tACS were applied at 20 Hz, with the latter consisting of only the fade-in and -out. For individualized tACS, frequency was personalized (19.0 ± 3.0 Hz, **Figure 1G**, **Appendix 3**), based on the β peak frequency in terms of power during baseline motor planning [27]. Frequencies were rounded to the nearest 0.5 Hz to comply with hardware limitations.

### 2.4. Electroencephalography

EEG data were recorded using a 64-channel BioSemi ActiveTwo EEG system, with caps fitted based on head size. Signal Gel was used and offsets were kept below |20| µV. Data were sampled at 2048 Hz (2021a). Preprocessing, done using EEGLAB and custom functions (cf., **Appendix 3** for the full pipeline), involved down-sampling to 512 Hz, low-pass filtering, bad channels removal and interpolation, and cleaning tACS artefacts via sine-wave fitting and subtraction, and Signal-Space Projection (SSP) [4]. SSP was applied to all the data to prevent spatial distortions. Data were visually cleaned, re-referenced to the average, and individual component analysis was used to remove bad components. Data were epoched (−2.65 to 5.3 s, 0 s = movement onset), with on average 35.6 ± 4.4 (block 1), 69.7 ± 10.6 (block 2), 36.6 ± 3.5 (block 3) epochs per block. Two datasets were excluded due to insufficient epochs (<20). Time-frequency decomposition (1 – 35 Hz in 1 Hz steps) (**Appendix 3**) was done using complex Morlet Wavelets per participant, session and block. Power was calculated as the squared sum of the real and imaginary components.

### 2.5. Statistical analyses

All models were constructed via stepwise backward building, systematically removing non-significant effects. Tukey-corrected post-hoc comparisons were done when applicable. Analyses were performed in MATLAB and Rstudio [37, 38]. Linear mixed effect models (LME) are reported without general- and subject-specific intercepts and error terms for conciseness, although these were present in all models. Alpha was set at 0.05, and all P-values were two-tailed.

#### 2.5.1. The effect of β-tACS on β-band desynchronization

The effects of tACS on MRβD were examined in threefold (**Figure 2**). All analyses focused on EEG activity at the tACS frequency ± 2 Hz. For fixed and sham tACS, this was 20 Hz. For individualized tACS, frequency was personalized.

**Figure 2.**
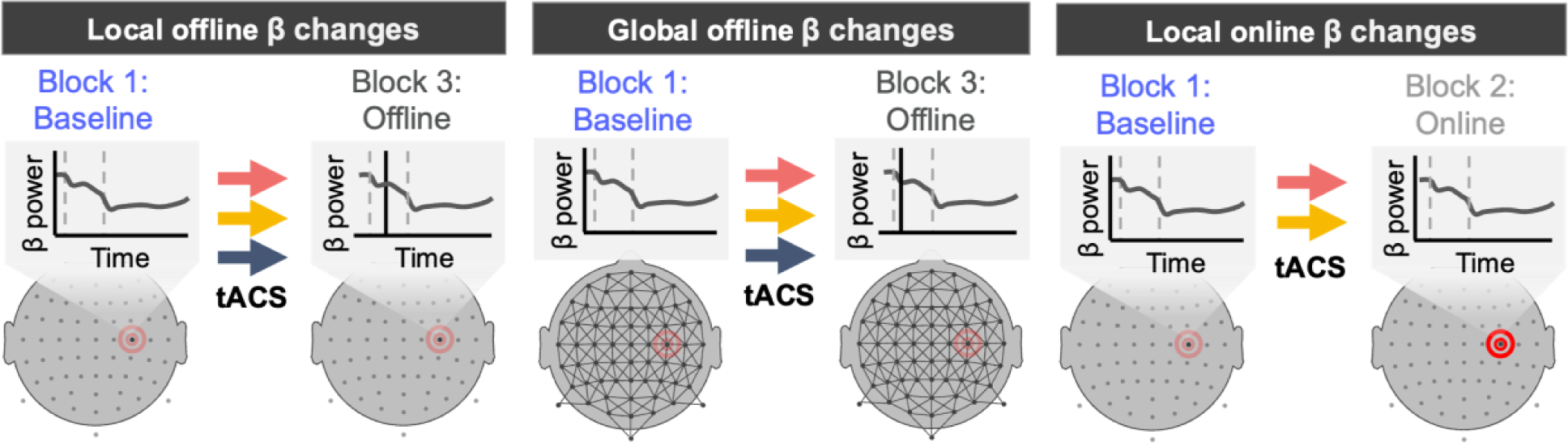
The threefold analysis of effects on the tACS frequency band ±2 Hz. Colored arrows indicate fixed (red), individualized (yellow) and sham tACS (blue). The red circles on the topographic plots represent the tACS montage. Left: A first analysis gauged local offline effects. Middle: A second analysis gauged broad effects using a spatiotemporal clustering approach, with black lines representing spatial adjacency. Right: The third analysis examined local online effects.

The first two analyses assessed local and global offline effects, while the third analysis investigated local online effects. Online and offline EEG data were separately analyzed due to tACS artefact removal challenges. While our artefact removal pipeline produced similar neural signatures for the offline and online data, separating the data eliminated the risk of tACS-artefacts affecting conclusions drawn based on the offline data, while still providing the potential of novel insights into the immediate neurophysiological effects of tACS.

##### 2.5.1.1. Local MRβD changes at the tACS frequency

This analysis focused on local changes from baseline to the offline block. We applied a temporal clustering approach, fitting an LME per timepoint from 0 to 4 s:

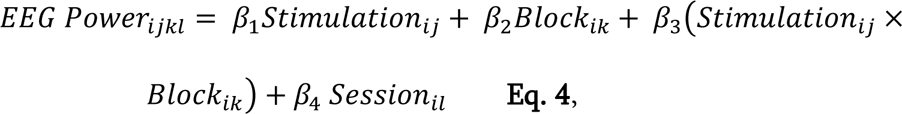

with EEG power relating to the mean power in the tACS stimulation frequency ± 2 Hz. F-values were retained and used to calculate threshold-free clustering enhanced (TFCE) values [39]:

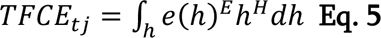

where *h* is cluster height (F-value cut-off), *e* is cluster extent (the number of temporally adjacent datapoints > *h)*, and *E* and *H* are their respective weights, defaulting to *E* = 0.5 and *H* = 2. Starting at 0, *h* increased in 0.2 steps until the maximum F-value was reached. Significance was inferred if original TFCE values exceeded the 95^th^ percentile of a surrogate null distribution generated from 800 permutations.

##### 2.5.1.2. Global MRβD changes at the tACS frequency

This analysis explored global offline changes through spatiotemporal clustering. Mean EEG power per timepoint and sensor was fitted via an LME (cf., **Equation 4**), and the resulting F-values were used to calculate TFCE values (cf., **Equation 5**) [39]. Adjacency was not only temporal, but also spatial, based on neighboring sensors (**Figure 2**, middle panel). Significance inference was identical to **Section 2.5.1.1**.

##### 2.5.1.3. Online effects of tACS on MRβD

This analysis examined whether fixed and individualized tACS have different online effects. The sham condition was excluded due to the attenuation of the EEG signal by the tACS-artefact removal pipeline (**Figure 3**).

**Figure 3.**
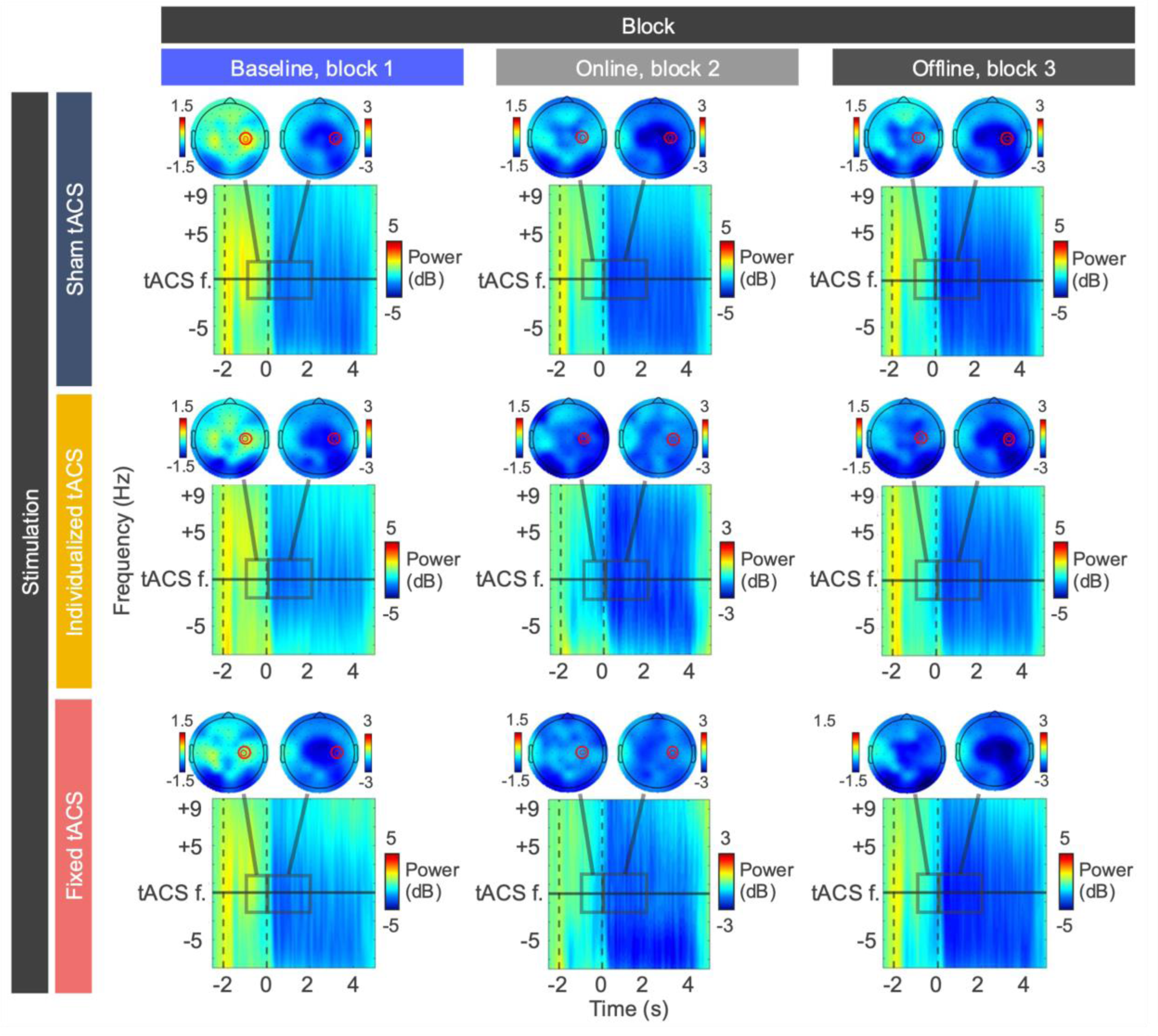
C4 time-frequency matrices and topographic plots representing MRβD changes across blocks and conditions. Time is locked to movement onset (0 s). Frequency is scaled based on the tACS frequency, which was either individualized or 20 Hz. Topographic plots show mean activity at the tACS frequency ±2 Hz in the final second of motor planning and initial 2 seconds of execution.

Per timepoint, we calculated the change in EEG power by subtracting the baseline block mean from the online block mean. An LME was used to test the effect of stimulation on the change in EEG power per timepoint:

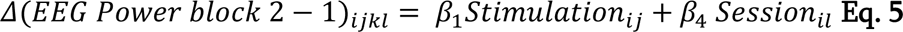

Temporal clustering was again used and significance was inferred in line with **Section 2.5.1.1**.

#### 2.5.2. The effect of β-tACS on bimanual motor performance

We first assessed whether baseline TE was similar across the tACS conditions via an LME, with TE as dependent variable and Stimulation as independent variable.

We then assessed the behavioral effects of tACS via temporal clustering, acknowledging the BTT as time-series data. An LME was fitted per timepoint;

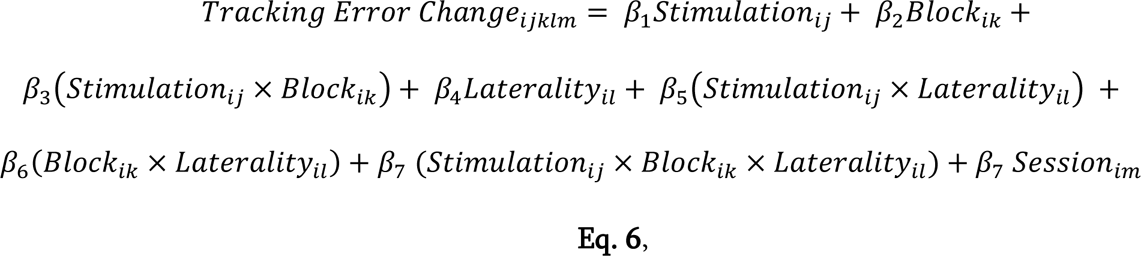

with laterality referring to whether both hands moved at the same speed (‘iso’), or faster rotations were required with the left or right hand. The TFCE procedure and permutation testing were identical to **Section 2.5.1.1**. In **Appendix 4**, we examined tACS effects on mean TE Change per trial instead of per timepoint. This resembles how previous research analyzed the BTT.

#### 2.5.3. The link between β-band activity and bimanual motor performance

Our final analysis examined whether MRβD changes during motor planning and/or execution were related to motor control changes, providing mechanistic insights into the polymorphic nature of MRβD. We extracted the 10^th^ percentile MRβD magnitude in C4 during motor planning and execution. The difference in peak MRβD from the offline block to baseline was computed, and an LME was fitted:

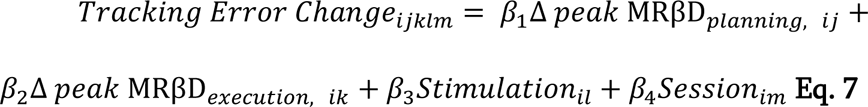

#### 2.5.4. Assessing tACS blinding

Blinding effectiveness was assessed using the Chi-Squared test to determine whether participants’ responses to the question of whether they believed they received verum, sham or were unsure, were independent of the administered tACS condition [40].

## 3. Results

### 3.1. Study Sample

**Table 1** outlines general sample characteristics. In total, 8% of the EEG data and 7.4% of the BTT data were excluded due to tACS bridging (6.5%) –identified by unstable and impedance values below 1 kΩ–, EEG data removal (0.6%) due to too few epochs, and study-unrelated illness resulting in a drop-out after session 2 (0.9%). No tACS side-effects occurred and all participants tolerated tACS well.

**Table 1.**
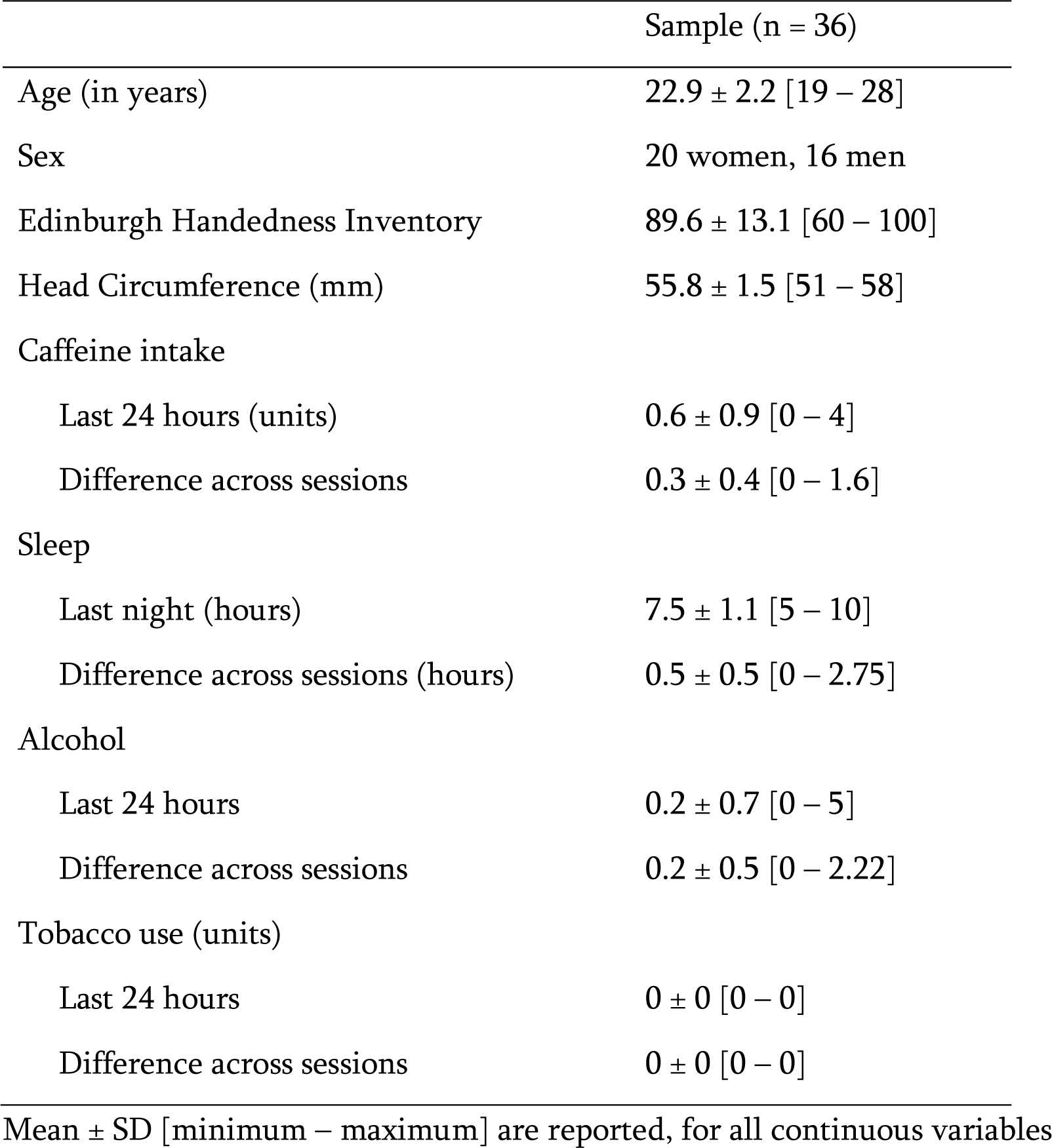
Sample characteristics.

|  | Sample (n = 36) |
| --- | --- |
| Age (in years) | 22.9 ± 2.2 [19 – 28] |
| Sex | 20 women, 16 men |
| Edinburgh Handedness Inventory | 89.6 ± 13.1 [60 – 100] |
| Head Circumference (mm) | 55.8 ± 1.5 [51 – 58] |
| Caffeine intake |  |
| Last 24 hours (units) | 0.6 ± 0.9 [0 – 4] |
| Difference across sessions | 0.3 ± 0.4 [0 – 1.6] |
| Sleep |  |
| Last night (hours) | 7.5 ± 1.1 [5 – 10] |
| Difference across sessions (hours) | 0.5 ± 0.5 [0 – 2.75] |
| Alcohol |  |
| Last 24 hours | 0.2 ± 0.7 [0 – 5] |
| Difference across sessions | 0.2 ± 0.5 [0 – 2.22] |
| Tobacco use (units) |  |
| Last 24 hours | 0 ± 0 [0 – 0] |
| Difference across sessions | 0 ± 0 [0 – 0] |

Mean ± SD [minimum – maximum] are reported, for all continuous variables

### 3.2. The effect of β-tACS on MRβD

**Figure 3** shows MRβD across blocks and conditions. MRβD during motor execution was consistent across blocks, while it emerged during motor planning from block 2 onward.

#### 3.2.1. Local MRβD increases as a result of block and session

A temporal clustering analysis examined tACS effects on MRβD (**Figure 4A**). Our findings concerning tACS were in the direction of our hypothesis, as mean MRβD during motor planning was enhanced in the fixed and individualized tACS conditions compared to sham. However, both this effect and the stimulation*block interaction were insignificant, as the TFCE value was below the 95^th^ percentile of the null distribution.

**Figure 4.**
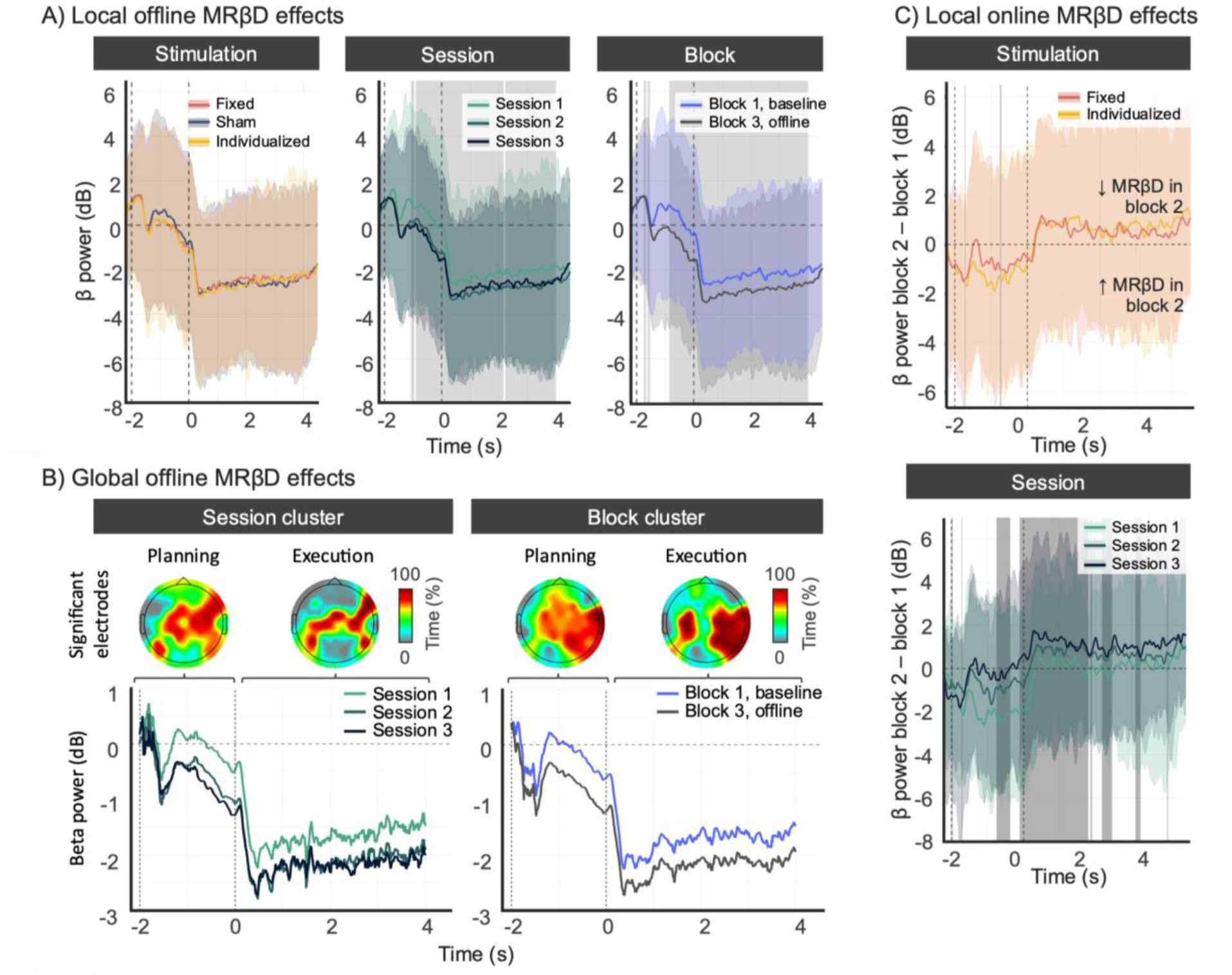
Effect of tACS on MRβD. **A)** Local offline effects of stimulation (left), session (middle), and block (right) on MRβD in the target sensor, C4. Grey areas denote significance. There was no significant effect of stimulation type. Conversely, session and block affected MRβD, with increases over sessions and blocks. **B)** Global offline effects of session (left) and block (right) on MRβD, the upper plots show the percentage of time during which data in sensors significantly differed across conditions (sessions and blocks) during motor planning and execution. The lower plots show MRβD time course within the sensors with significantly different activity. Whole-sensor session and block effects resembled the local effects discussed in A). **C)** Local online effects of stimulation (upper) and session (lower) on MRβD in the target sensor, C4. Grey areas denote significance. More negative values indicate a greater increase in MRβD in the online block compared to baseline. Concerning stimulation, two transient clusters were significant, with MRβD during motor planning being slightly more enhanced in individualized vs. fixed tACS. The TFCE values associated to these clusters are also shown in Appendix 5. Concerning session, MRβD in the online block vs. baseline increased the most in session 1, and least in session 3.

Session affected MRβD, with the highest MRβD during both motor planning and execution in session 3, and the lowest in session 1, and differences between sessions 2 and 3 being minor. Three clusters were significant: a transient first cluster (−1072 to −963 ms, peak F_2,191_ = 4.120), followed by longer-lasting clusters 2 (−908 to 2147 ms, peak F_2,191_ = 17.401) and 3 (2225 to 4000 ms, peak F_2,191_ = 7.204).

MRβD increased from baseline to the offline block, during both motor planning and execution. Three clusters were significant: cluster 1 (−1752 to −1674 ms, peak F_1,191_ = 9.753 at −1705 ms), 2 (−1619 to −1533 ms, peak F_1,191_ = 10.168 at −1580 ms), and 3 (−861 ms to 4000 ms, F_1,191_ = 46.19 at 490 ms).

#### 3.2.2. Broad MRβD increases as a result of block and session

Beyond local effects, we analyzed whole-sensor effects through spatiotemporal clustering. No tACS effects or interactions survived multiple comparison correction. Conversely, session and block significantly affected MRβD (**Figure 4B**).

MRβD magnitude increased with sessions in the significant spatiotemporal sensors (**Figure 4B**). The effect was more spatially pronounced during motor planning (**Figure 4B**, upper left plot), particularly in the right frontocentral, central, centroparietal, and left medial-central sensors. During execution, sensors containing significant effects were mainly restricted to the left and right central sensors (**Figure 4B**).

MRβD magnitude also increased from baseline to the offline block, with a more spatially extensive effect during motor planning compared to execution, particularly in the midline and right frontocentral, central and centroparietal sensors (**Figure 4B**). Significant effects during execution were present in the left and right centroparietal sensors.

#### 3.2.3. Stimulation and session affect the changes in MRβD from block 1 (baseline) to block 2 (online)

We also examined if MRβD changes from baseline to the online block were influenced by tACS and session. This analysis focused only on the verum tACS conditions due to the potential impact of the artefact removal pipeline.

Overall, MRβD changes during motor planning from baseline to the online block were most pronounced for individualized tACS (**Figure 4C**), with two significant transient clusters: cluster 1 (−1720 to −1705 ms; peak F_1,60_ = 6.0835) and 2 (−751 to −721 ms; peak F_1, 60_ = 5.703). The TFCE full-time course of this effect is shown in **Appendix 5**. No significant stimulation effects were present during motor execution, with both groups showing no-to-marginal decreases in online MRβD compared to baseline.

Session also affected MRβD, with the most substantial enhancements from baseline to block 2 in session 1, followed by sessions 2 and 3. The pronounced changes in session 1 align with greater BTT improvements in this session (**Section 3.3**), supporting the hypothesis that MRβD during motor planning represents processes relevant to motor control. Eight significant session clusters were present (**Figure 4C**, lower panel), the three largest being clusters 1 (−752 to −369 ms; peak F_2,60_ = 7.836), 2 (111 to 1787 ms; peak F_2,60_ = 12.027) and 3 (2170 to 2451 ms, peak F_2,60_ = 9.573).

### 3.3. Individualized β-tACS improves bimanual motor control

Baseline BTT performance was similar across conditions, as the LME found no significant effect of tACS condition (F_2, 268.42_ = 0.889, p = 0.41).

Stimulation affected TE change, with a significant cluster from 0.6 to 4 s (peak F_2, 2342_ = 18.721) (**Figure 5**). The greatest improvement in BTT performance was present in the individualized tACS condition (estimate = −0.53 ± 0.70), followed by sham (estimate = −0.46 ± 0.66) and fixed tACS (estimate = −0.40 ± 0.62). The traditional, mean-trial, analysis (cf., **Appendix 4**) corroborates this, with Tukey-corrected post-hoc tests showing significant differences between individualized and fixed tACS, and individualized and sham tACS.

**Figure 5.**
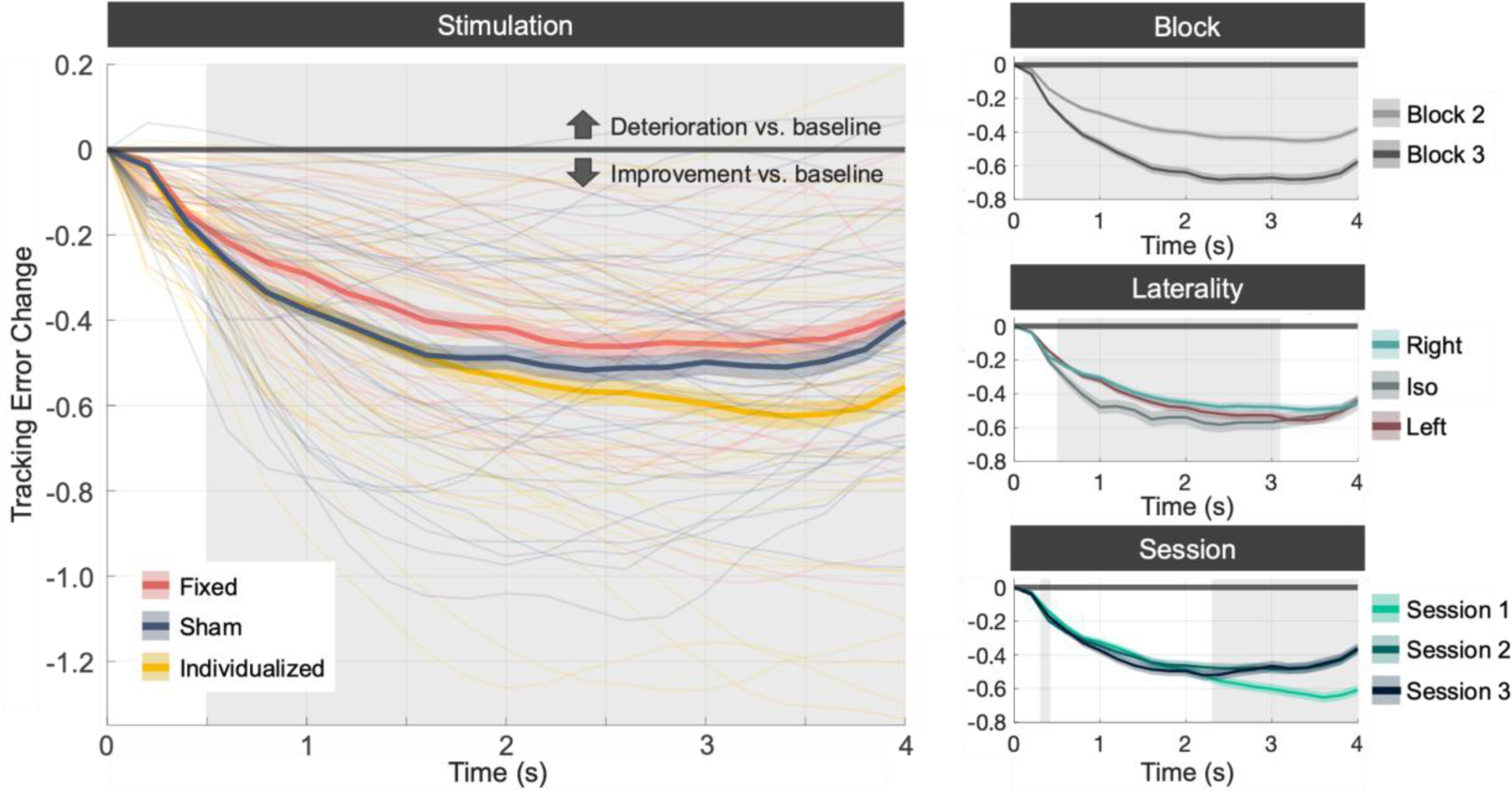
Effect of stimulation, block, laterality and session on tracking error change per timepoint of the BTT task. Tracking error change denotes the change in tracking error relative to the baseline block. Grey areas denote significant timepoints (p < 0.05), colored areas denote 95% confidence intervals.

The significant effect of block (peak F_1, 2342_ = 91.126) spanned from 0.2 to 4 s, with the improvement in TE with respect to block 1 being greatest in block 3 (estimate = −0.57 ± 0.73), compared to block 2 (estimate = −0.36 ± 0.57).

Concerning laterality (peak F_2, 2342_ = 31.192, p < 0.05), a cluster from 0.6 to 3 s was present. Improvement in this cluster was largest for the iso (estimate = −0.56 ± 0.86), left (estimate = −0.47 ± 0.54) and right conditions (estimate = −0.44 ± 0.51).

Concerning session (peak F_2, 2342_ = 54.930, p < 0.05), a cluster from 2.2 to 4 s was retained. BTT improvement was greatest in session 1 (estimate = −0.70 ± 0.72), followed by session 2 (estimate = −0.50 ± 0.72) and 3 (estimate = −0.48 ± 0.73).

A post-hoc LME (TE Change ∼ tACS FREQUENCY * BLOCK * LATERALITY + SESSION) found no significant interaction or main effects of tACS frequency. This suggests that the effectiveness of individualized tACS was not due to stimulation at a fixed frequency that happened to be more effective than 20 Hz, but rather due to its individualized nature.

**Figure 6.**
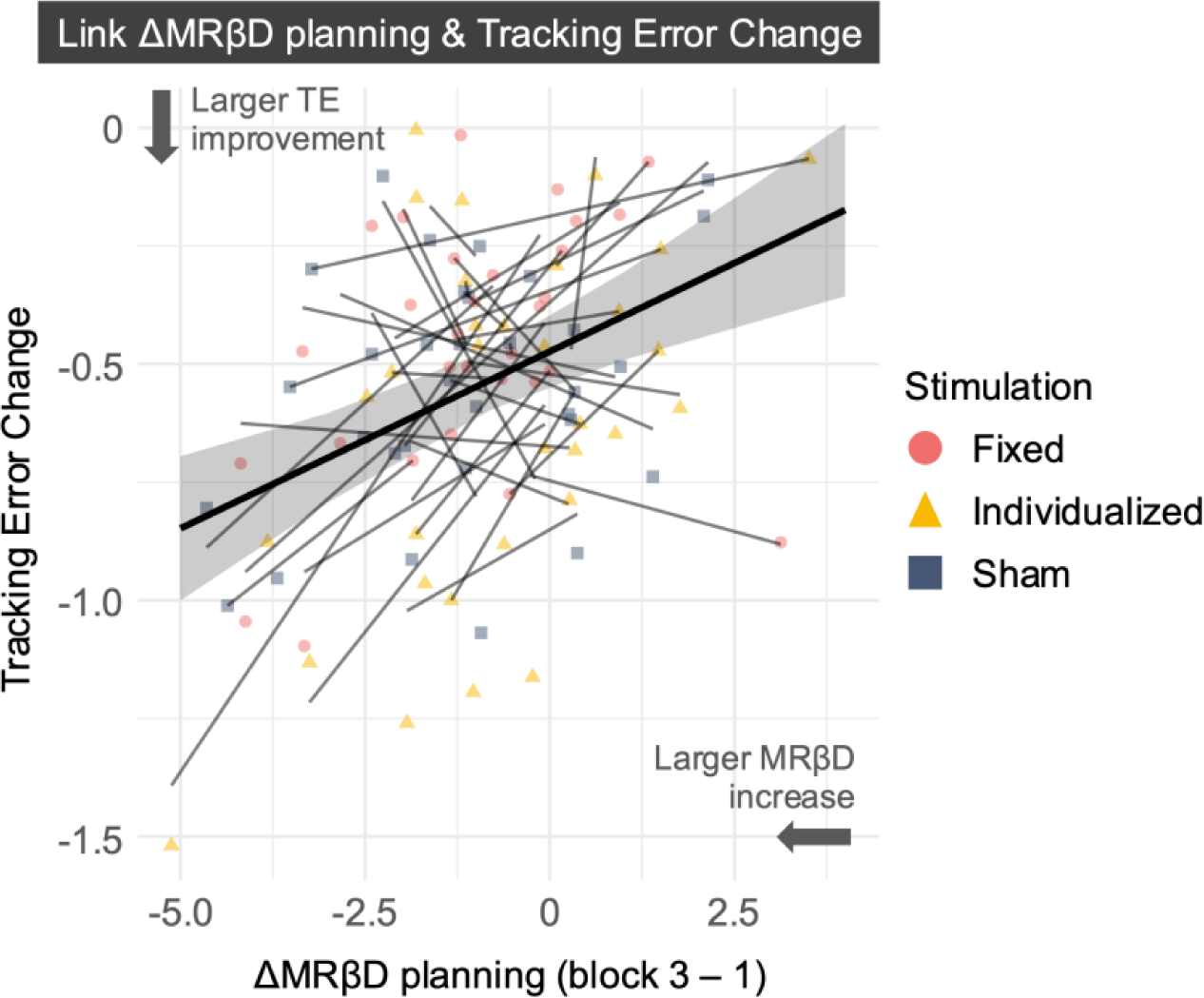
The relationship between change in tracking error (TE) and motor planning MRβD from baseline to the offline block. Larger MRβD increases were associated to larger bimanual tracking task improvements.

### 3.4. The link between β-band desynchronization and BTT performance

We investigated if changes in motor planning and/or execution MRβD from baseline to the offline block were related to TE changes, when controlling for the effects of stimulation and session.

Consistent with our hypotheses, we found a significant effect of change in MRβD during motor planning (F_1, 86.088_ = 18.721, p < 0.001) but not during motor execution. Increased MRβD during motor planning from baseline to the offline block was associated to larger BTT improvements from baseline to the offline block. The covariate stimulation was also significant (F_2, 62.501_ = 4.057, p = 0.022), as discussed in **Section 3.3**.

### 3.5. tACS blinding effectiveness was limited

There was a significant association between STIMULATION TYPE and tACS beliefs (χ²_4_ = 14.391, p = 0.006), with corrected comparisons revealing differences between fixed- and sham tACS (p = 0.045) and individualized and sham tACS (p = 0.016), but not between both verum tACS conditions (p < 0.05). While the number of participants believing that they received verum tACS was similar in fixed (58%) and individualized (63%) tACS, it was lower for sham (31%). Conversely, identification as sham was highest in sham (41%) compared to fixed (12%) and individualized tACS (9%). The number of participants uncertain whether they received verum or sham was similar across conditions: 30% for fixed tACS, 28% for individualized tACS, and 28% for sham.

## 4. Discussion

The current work applied individualized-, fixed 20 Hz- and sham tACS in 36 healthy adults across three days while performing a continuous bimanual motor task. Our main findings are that individualized β-tACS improves bimanual motor performance, that MRβD magnitude is positively associated to bimanual motor control, and that individualized β-tACS seems to enhance online MRβD. Together, these results underscore the importance of personalizing tACS and the relevance of MRβD in motor control.

### 4.1. The Functionally Polymorphic Nature of MRβD in Motor Control

MRβD has been described as functionally polymorphic, playing distinct roles during motor planning and execution. In motor planning, MRβD may reflect processes of somatosensory integration and preparation of motor commands [23, 28, 41], influenced by factors like motor complexity in interlimb tasks [27, 28, 42], directional uncertainty during reaching [25], and effector involvement in grasping [43]. Conversely, during motor execution, MRβD appears to represent more general motor processes, with findings suggesting insensitivity to motor complexity [27], the speed-accuracy trade-off [44], and grasp type [45]. Here, we build on the hypothesis of MRβD’s functional polymorphism, as we found a positive association between MRβD and bimanual motor control during motor planning, but not execution. Moreover, MRβD during motor planning increased as participants became more proficient within and across sessions. The observed MRβD increase during motor execution seems to be driven by heightened MRβD during planning, as illustrated in **Figure 4**. This concurs with our previous work, where increased MRβD during motor planning seemed to drive increased MRβD during execution [27].

### 4.2. Individualized tACS Enhances Bimanual Motor Control And Online MRβD

Research into the online effects of tACS on event-related perturbations is limited, primarily due to associated artefacts, especially at intensities such as 3 mA peak-to-peak. The few available studies indicate that tACS at lower intensities (around 1.5 mA) enhances event-related perturbations [17–19].

We demonstrate that individualized tACS increases online MRβD during motor planning compared to fixed tACS. While a similar trend was observed offline for both individualized and fixed tACS, it was not statistically significant. Notably, individualized tACS improved bimanual motor control compared to sham and fixed tACS. This improvement was not explained by the absolute tACS frequency, supporting the idea that individualized tACS is more effective due to better alignment with participants’ endogenous brain activity, consistent with the Arnold Tongue principle [8].

A theoretical framework for these findings combines said Arnold Tongue principle with recent computational models, which suggest that low-intensity tACS initially desynchronizes neural firing by disrupting endogenous synchronization, and only starts to induce net synchronization when intensity increases [12]. Given that we applied tACS at a low intensity (cf., **Figure 1D** and [12]), we may have caused this desynchronization effect. However, due to the baseline correction needed for artifact removal [21], it is impossible to untangle if our observed MRβD increases resulted from tACS desynchronization (cf., above) or merely reflect a need to cope with increased baseline β-activity as a result of tACS-induced entrainment. More invasive neuroimaging methods which can achieve higher signal-to-noise ratios should investigate this further.

Both tACS conditions appeared to slightly attenuate online MRβD during motor execution (**Figure 4C**). As this effect occurred in both conditions, it is likely a result of the artifact removal pipeline, as SSP is known to attenuate neural signals with topographies similar to tACS artifacts [4]. If so, this may suggest that the MRβD enhancement during motor planning could be stronger than indicated by the current results or that its topography differed from MRβD during motor execution and the tACS artefact, resulting in less pronounced attenuation due to SSP. Previous research showing distinct β-activity topographies before and after movement provides an argument in favor of the latter [28].

The lack of significant offline tACS effects on MRβD may reflect the transient nature of tACS [46]. However, offline neuroplasticity-like tACS effects are being increasingly recognized, with previous work showing an offline enhancement of α power following individualized tACS [5, 11, 16]. Such effects may have occurred in the current study, given that the behavioral effects of individualized tACS did persist, although they would imply that the MRβD did not represent them. Alternatively, data variability and/or the subtlety of post-stimulation effects may have limited detection of such effects. Given that mean offline MRβD trended in the hypothesized direction, this remains a plausible explanation.

### 4.3. Hemispheric Asymmetry in MRβD

Although we did not directly compare tACS effects between hemispheres, our results concur with the notion that right sensorimotor regions are critical in complex motor control [27, 36, 47, 48]. While the left hemisphere is typically dominant from a movement point-of-view, the right hemisphere provides additional support when complexity increases [27, 28, 34, 36]. When comparing MRβD across hemispheres, right sensorimotor MRβD predicted motor control improvements best (**Appendix 6**). Likewise, for the global effects of block and session (**Section 3.2.2**.), particularly during motor planning, data related to the right sensors were predominantly significant. These observations support the importance of the right sensorimotor regions in complex motor behaviors such as bimanual motor control. Consequently, these findings imply that while most β-tACS montages have focused on the left sensorimotor cortex [16, 49], the right sensorimotor cortex may be a viable future target.

### 4.4. Limitations

Several limitations should be considered. Applying tACS at 3 mA peak-to-peak, in line with [16, 50], posed challenges in removing tACS artifacts, as the intensity was higher than what is typically used in concurrent tACS-EEG studies. While we recovered genuine neural activity –the online data showed a similar session effect as the offline data, and a tACS effect consistent with our hypotheses–, the online data required more interpolation of channels of interest and was attenuated resulting in the need to separately analyze the offline and online data. Also, the lack of an online sham condition limits our interpretation of online tACS effects. A significant portion of participants correctly identified the type of tACS they received.

This raises concerns about the effectiveness of blinding in tACS studies. While this does not affect the comparisons between fixed versus individualized tACS conditions, which is reassuring for our results, it implies that improvements in conventional sham protocols are needed. Shunting-based sham methods may enhance blinding and strengthen the rigor of future work [51].

Lastly, only participants were blinded due to the overtness of artefacts in the EEG data, and the need to individually set-up the tACS parameters in the individualized tACS condition.

## 5. Conclusion

Individualized β-tACS enhances bimanual motor control, with significant behavioral improvements highlighting the value of personalized neuromodulation. Although the effects on MRβD were subtle and transient, the strong link between MRβD during motor planning and motor task improvements underscores its critical, polymorphic role in motor control. These findings pave the way for individualized tACS in neurorehabilitation, indicate that the right sensorimotor cortex is an interesting tACS target in light of motor control in right-handed individuals, and offer a framework for studying the role of event-related perturbations in neural oscillations in motor control and beyond.

## CRediT authorship contribution statement

**SVH:** Conceptualization, Methodology, Software, Validation, Formal Analysis, Investigation, Resources, Data Curation, Writing – Original Draft, Writing – Review & Editing, Visualization, Project Administration, Funding Acquisition.

**DABM:** Conceptualization, Methodology, Software, Validation, Investigation, Writing – Review & Editing, Supervision.

**MN:** Conceptualization, Writing – Review & Editing

**KC:** Writing – Review & Editing, Resources, Funding Acquisition.

**SV:** Conceptualization, Writing – Review & Editing, Supervision, Project Administration, Funding Acquisition.

**RF:** Conceptualization, Writing – Review & Editing, Supervision, Project Administration, Funding Acquisition.

## Acknowledgments

The authors wish to thank Ing. Marc Geraerts for technical support, and the many master students of Rehabilitation Sciences and Physiotherapy who assisted in data acquisition.

## Declaration of competing interest

The authors report no conflicts of interest.

## Funding

SVH was funded by a FWO PhD fellowship (G1129923N) and additional BOF bench fee (BOF22INCENT19). DABM was funded by research visiting funds granted by the ‘Korte verblijf programma’ (BOF23KV15). MN was funded by a FWO PhD fellowship (11PBG24N) and an additional BOF bench fee (BOF23INCENT18).

## Appendices

## A1. Hard- and software manufacturer information

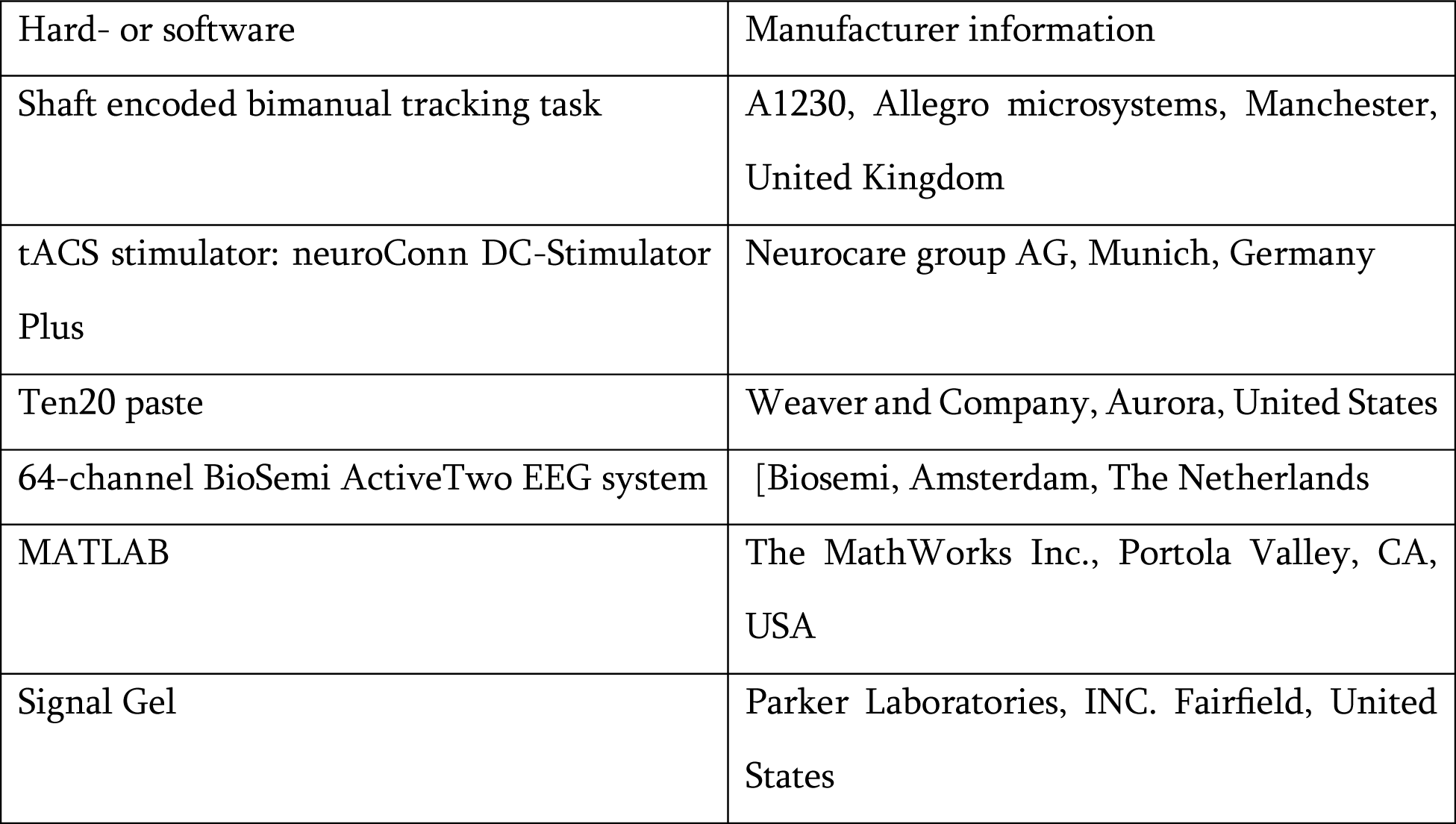

## A2. In- and Exclusion criteria

Via G*Power (v.3.1), the estimated sample size for a power of 0.80 was n = 24 (effect size f = 0.3, extracted from [52]). Considering drop-outs and loss-of-data, we instead aimed to recruit 36 healthy participants.

Prior to participation, the following criteria were assessed. Inclusion criteria were:

- Age between 18 and 30 years old
- Right-handed according to the Edinburgh Handedness Inventory [53]
- (Corrected-to-)normal sight
- Healthy, self-reported, and defined as the absence of any exclusion criteria

Exclusion criteria were:

- No neurological, psychiatric and/or cognitive disorders
- No metal implants in the head and/or spine
- No epilepsy
- No use of medication that affects the central nervous system
- No allergy to lotions and/or cosmetics
- No smoking
- No drug- and/or alcohol addiction
- No physical impairment making the performance of the BTT impossible
- No musical instrument use

## A3. EEG Preprocessing

### Pipeline for offline data analysis

Data were downsampled to 512 Hz, 1 Hz forward-backward low-pass filtered, and bad channels were identified via visual inspection (removed channels: 3.8 ± 2.9) and interpolated.

tACS-contaminated data were cleaned by first fitting and subtracting a sine-wave per channel and 2 s epoch. As this only suppresses stationary artefacts, Signal-Space Projection [4] was subsequently applied. Briefly, singular value decomposition was performed on 2 s epochs. The most prominent spatial pattern was projected out using a leadfield generated from the BioSemi cap lay-out and the MNI152 template MRI scan in Fieldtrip [54]. As this spatially distorts the signal, source-informed reconstruction was performed on all the EEG data. For more information, see [4].

Bad data portions were removed based on visual inspection, and data were re-referenced to the average reference. Individual component analysis was performed using Picard (maximum 500 iterations) and manually identified noisy components were removed (removed components: 3.6 ± 4.0). Epochs were made from −2.65 to 5.3 s, with 0 s being movement onset. An average of 35.6 ± 4.4 epochs remained for block 1, 69.7 ± 10.6 epochs for block 2, and 36.6 ± 3.5 epochs for block 3. Two datasets were excluded due to insufficient epochs (<20 epochs).

### EEG time-frequency decomposition

Per participant, session and block, the epoched data were decomposed into time-frequency representations by complex Morlet Wavelet convolution with complex Morlet wavelets, defined as Gaussian-windowed complex sine waves:

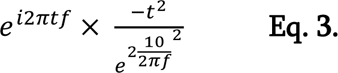

where *i, t,* and *f* are the complex operator, time and frequency. Time ranged from the epoch’s onset to its end. Frequency ranged from 1 to 35 Hz in 1 Hz increments. Power values were computed as the sum of the squared real and imaginary components per frequency and timepoint.

### Pipeline for online peak β extraction

The EEG data were downsampled to 128 Hz and 1 – 35 Hz forward-backward bandpass filtered. Next, bad channels and data sections were removed through clean_rawdata, followed by an interpolation of the removed channels. The data were re-referenced to the common average reference, and a rank-adjusted individual component analyses was performed. Subsequently, eye, brain, muscle, heart and line noise components were removed using ICLabel and the data were epoched from −2 to 0 s, with 0 s being movement onset. Lastly, the Fast Fourier Transform (FFT) was applied on the epoched data to acquire the frequency with the largest power within the β band (13.5 – 30 Hz), based on the Power Spectral Density. This frequency was withheld and used as tACS frequency for the individualized condition.

## A4. Traditional trial-wise analysis of BTT data

Next to the temporal clustering approach, we used a linear mixed model to examine tACS effects on BTT performance. While this does not acknowledge that the BTT results in time-series data, it is consistent with how previous work analyzed the BTT [27, 34].

A linear mixed model was constructed;

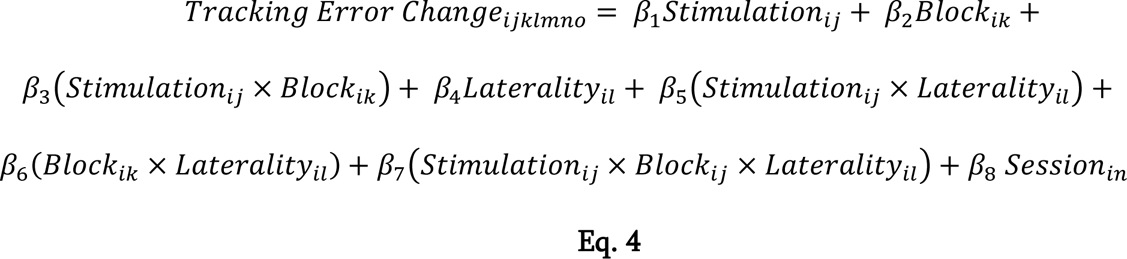

Where *β*_1_to *β*_7_ relate to the fixed effects of stimulation type, j, block, k, laterality, l, and their interactions, and the *β*_8_refers to the covariate for session, n. TE change values were aggregated to have one value per combination of stimulation, block, laterality, session and participant.

The final model, shown in **Table A4.1**., demonstrates significant effects of STIMULATION (F_2, 556.76_ = 11.305, p < 0.001), BLOCK (F_1, 544.35_ = 67.003, p < 0.001) and LATERALITY (F_2, 544.35_ = 4.213, p = 0.015), as well as for the covariate of SESSION (F_2, 556.80_ = 5.864, p = 0.003). No interaction effects remained after model building.

**Table A4.1.**
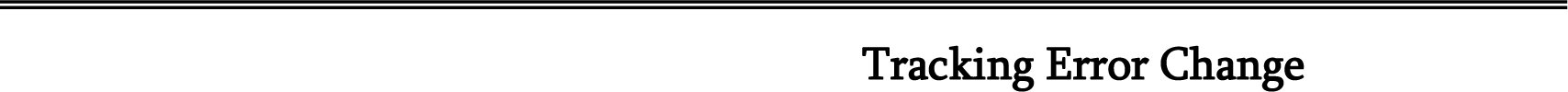
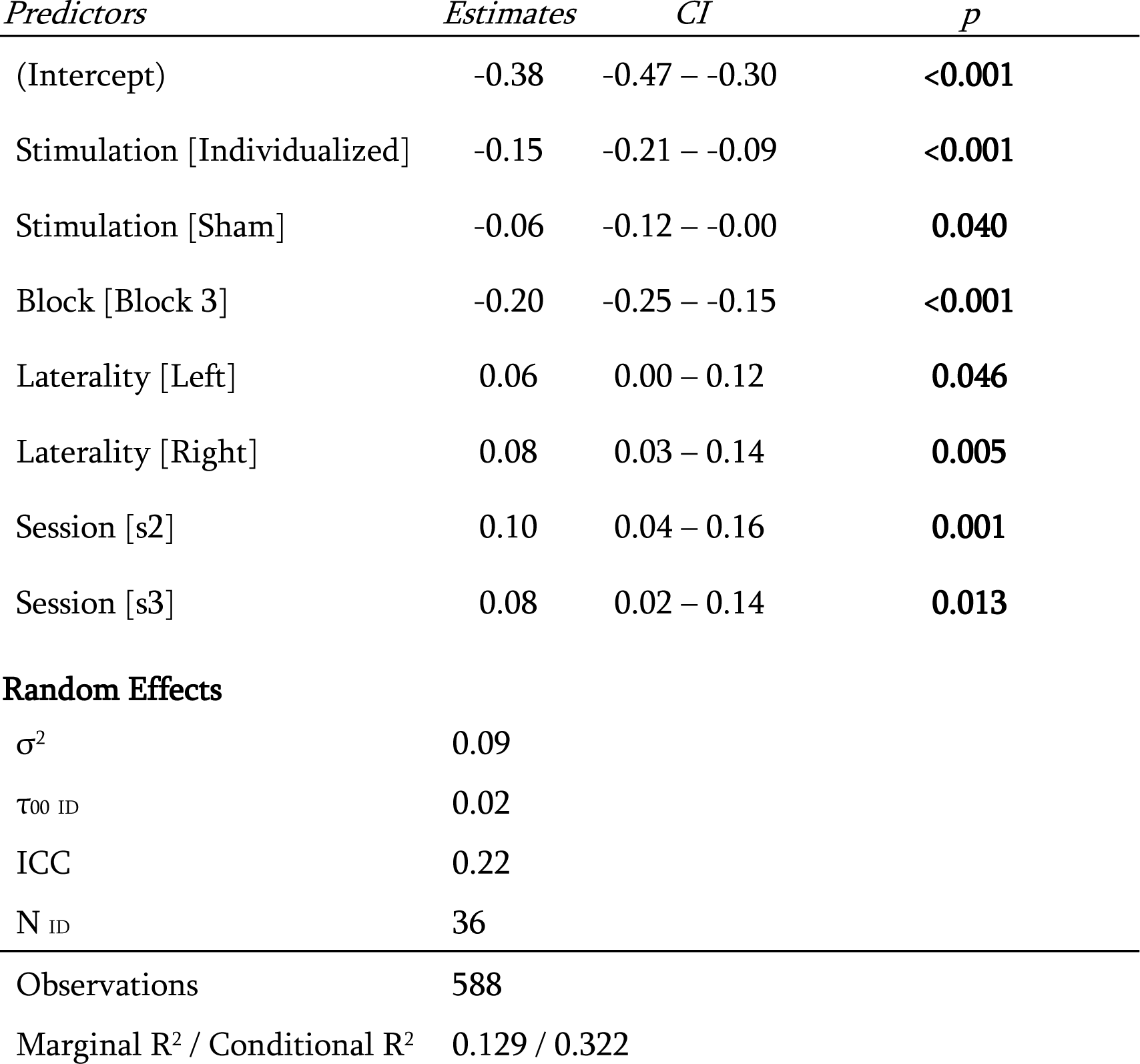
Linear Mixed Effect Model for tACS effect on tracking error change.

Tukey-corrected post-hoc tests for the effect of STIMULATION show that individualized tACS resulted in a greater improvement on the BTT than sham tACS (estimate = −0.09, t_555_ = −2.848, p = 0.013) and fixed tACS (estimate = −0.15, t_555_ = −4.727, p < 0.001). Concerning block (Table 2), as to be expected, BTT improvement was higher in block 3 than block 2. Post-hoc tests of the LATERALITY effect demonstrated that there was a significant difference between BTT conditions wherein both hands had to move at equal speeds, compared to conditions where the right hand had to move faster, with the latter conditions showing the greatest improvements with respect to block 1 (estimate = −0.08, t_545_ = −2.823, p = 0.013).

**Figure A4.1**. visually shows the main effects. As can be seen in this figure, there are some outliers with the most apparent ones being present in the individualized tACS condition. When rerunning the aforementioned model without outliers, defined by a distance larger than 3 standard deviations of the mean, the effect of STIMULATION and the associated post-hoc tests remained identical in terms of significance and interpretation.

**Figure A4.1.**
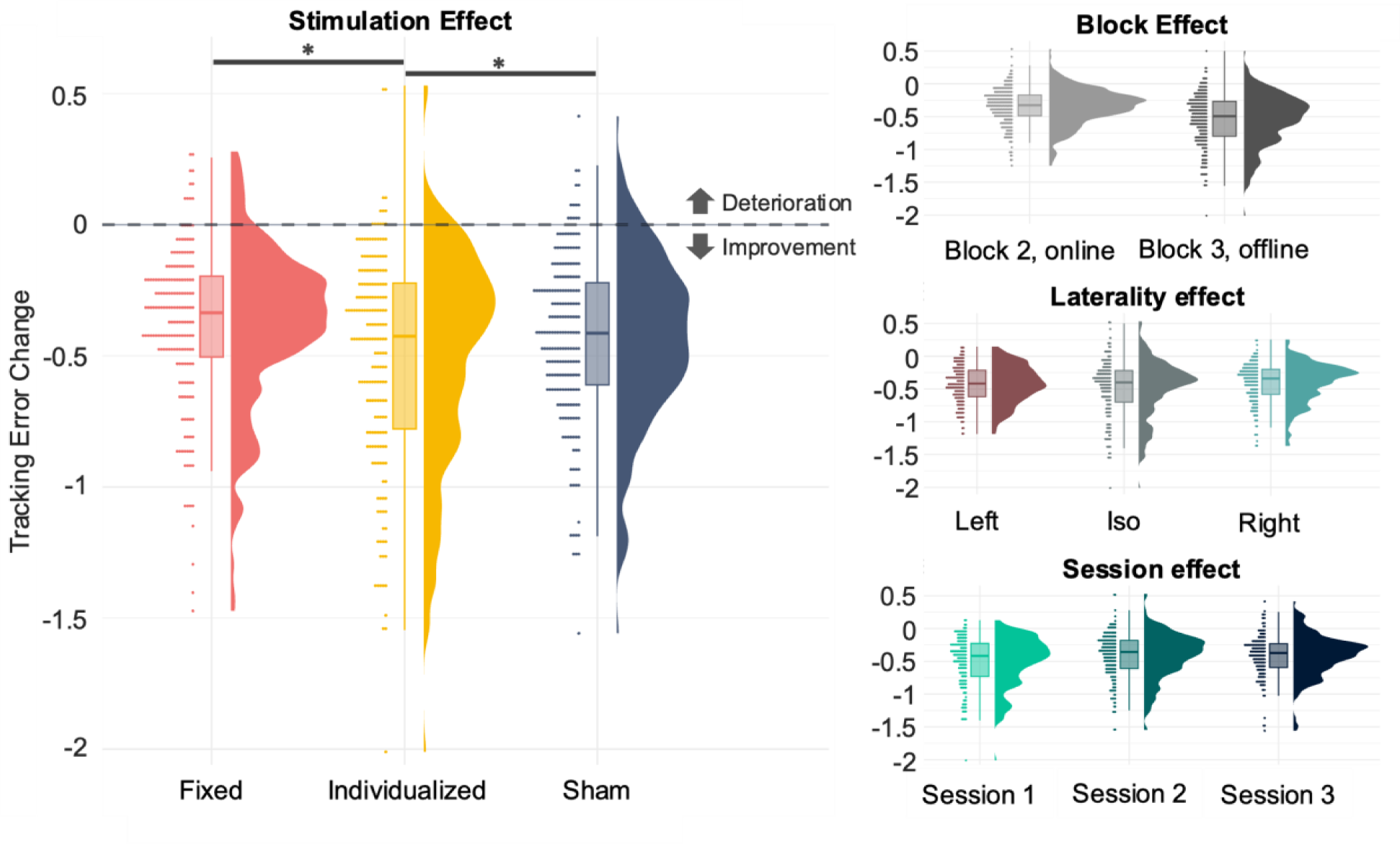
The effect of stimulation, block, laterality and session on mean tracking error change. The left panel shows the effect of stimulation, with greater increases in mean tracking error as a result of individualized tACS. The right upper panel shows the effect of block, with a larger tracking error improvement in block 3, compared to block 1.

## Appendix 5. Full Threshold Free Clustering Enhanced (TFCE) time course of online stimulation effect

**Figure A5.**
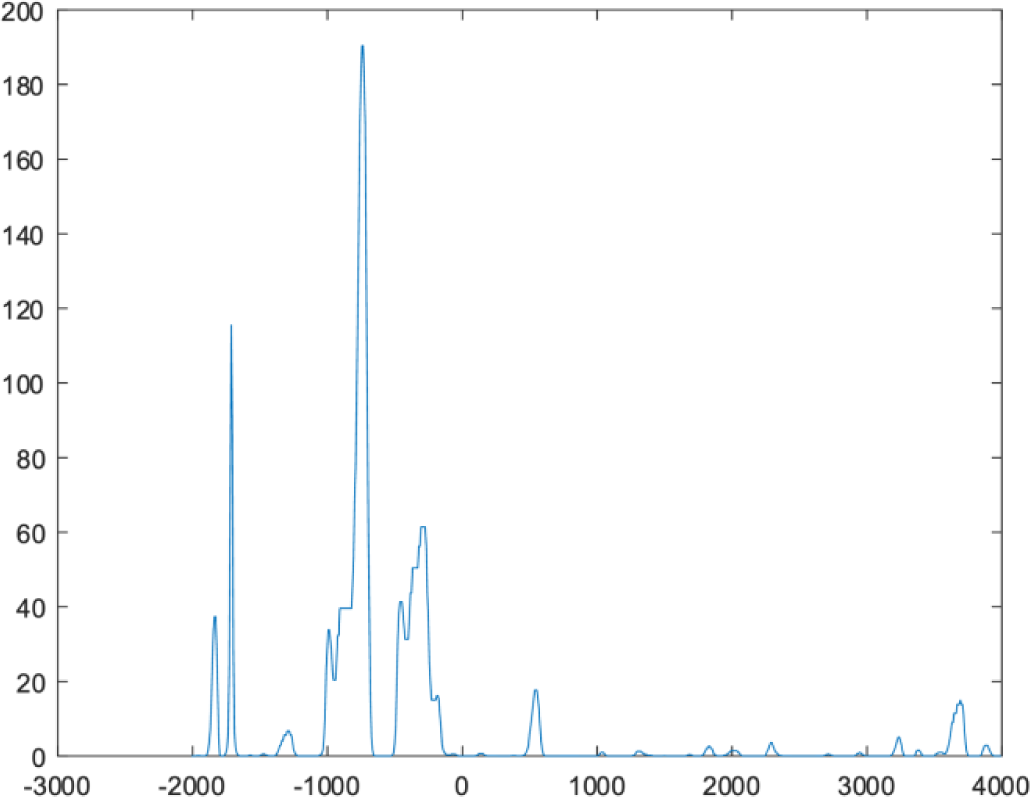
TFCE time course of the online stimulation effect on MRβD (fixed vs. individualized tACS). The Y axis denotes TFCE values, the X-axis denotes time in ms. As can be seen, the largest TFCE values occur during the planning phase (−2000 to 0 ms), whereas values TFCE during motor execution are low.

## A6. Comparison of the right and left EEG sensors to predict bimanual motor control improvements

Previous work suggests that the right sensorimotor cortex is particularly sensitive to increased motor task demands. To further test this with the available data, we aimed to see if perhaps data from EEG sensor C3 (left hemisphere) was better able to predict tracking error change than EEG sensor C4.

We ran the model defined in **Section 3.4**. using EEG data from sensors C3 or C4 and compared both models. The model used in the manuscript, incorporating sensor C4, best explained changes in tracking error, as shown in **Table A6**.

**Table A6.**
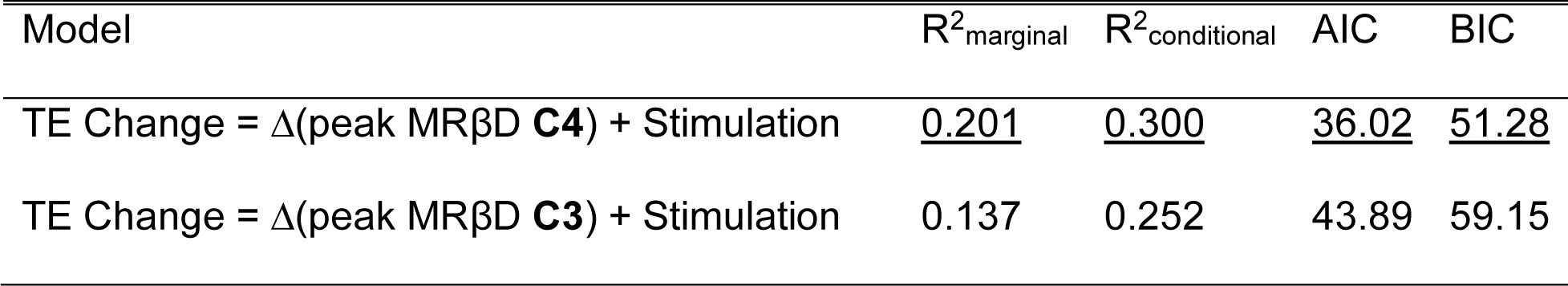
Comparison of TE Change prediction using peak planning MRβD extracted from the right (C4) and left (C3) sensorimotor regions.

| Model | R<sup>2</sup><sub>marginal</sub> | R<sup>2</sup><sub>conditional</sub> | AIC | BIC |
| --- | --- | --- | --- | --- |
| TE Change = $\Delta$(peak MRβD **C4**) + Stimulation | <u>0.201</u> | <u>0.300</u> | <u>36.02</u> | <u>51.28</u> |
| TE Change = $\Delta$(peak MRβD **C3**) + Stimulation | 0.137 | 0.252 | 43.89 | 59.15 |

## Notes

### Competing Interest Statement

The authors have declared no competing interest.

## References

1. Krakauer, J.W., Motor learning: its relevance to stroke recovery and neurorehabilitation. Curr Opin Neurol, 2006. 19(1): p. 84–90.

2. Nieuwboer, A., et al., Motor learning in Parkinson’s disease: limitations and potential for rehabilitation. Parkinsonism Relat Disord, 2009. 15 Suppl 3: p. S53–8.

3. Herrmann, C., et al., Transcranial alternating current stimulation: a review of the underlying mechanisms and modulation of cognitive processes. Frontiers in Human Neuroscience, 2013. 7(279).

4. Vosskuhl, J., et al., Signal-Space Projection Suppresses the tACS Artifact in EEG Recordings. Front Hum Neurosci, 2020. 14: p. 536070.

5. Wischnewski, M., I. Alekseichuk, and A. Opitz, Neurocognitive, physiological, and biophysical effects of transcranial alternating current stimulation. Trends Cogn Sci, 2023. 27(2): p. 189–205.

6. Fröhlich, F. and D.A. McCormick, Endogenous Electric Fields May Guide Neocortical Network Activity. Neuron, 2010. 67(1): p. 129–143.

7. Anastassiou, C.A., et al., Ephaptic coupling of cortical neurons. Nature Neuroscience, 2011. 14(2): p. 217–223.

8. Huang, W.A., et al., Transcranial alternating current stimulation entrains alpha oscillations by preferential phase synchronization of fast-spiking cortical neurons to stimulation waveform. Nature Communications, 2021. 12(1).

9. Zaehle, T., S. Rach, and C.S. Herrmann, Transcranial Alternating Current Stimulation Enhances Individual Alpha Activity in Human EEG. PLOS ONE, 2010. 5(11): p. e13766.

10. Vossen, A., J. Gross, and G. Thut, Alpha Power Increase After Transcranial Alternating Current Stimulation at Alpha Frequency (α-tACS) Reflects Plastic Changes Rather Than Entrainment. Brain Stimulation, 2015. 8(3): p. 499–508.

11. Kasten, F.H., J. Dowsett, and C.S. Herrmann, Sustained Aftereffect of α-tACS Lasts Up to 70 min after Stimulation. Frontiers in Human Neuroscience, 2016. 10.

12. Zhao, Z., et al., Intensity- and frequency-specific effects of transcranial alternating current stimulation are explained by network dynamics. Journal of Neural Engineering, 2024. 21(2): p. 026024.

13. Krause, V., et al., Beta Band Transcranial Alternating (tACS) and Direct Current Stimulation (tDCS) Applied After Initial Learning Facilitate Retrieval of a Motor Sequence. Frontiers in Behavioral Neuroscience, 2016. 10.

14. Pollok, B., A.-C. Boysen, and V. Krause, The effect of transcranial alternating current stimulation (tACS) at alpha and beta frequency on motor learning. Behavioural Brain Research, 2015. 293: p. 234–240.

15. Heise, K.F., et al., Distinct online and offline effects of alpha and beta transcranial alternating current stimulation (tACS) on continuous bimanual performance and task-set switching. Scientific Reports, 2019. 9.

16. Wischnewski, M., D.J.L.G. Schutter, and M.A. Nitsche, Effects of beta-tACS on corticospinal excitability: A meta-analysis. Brain Stimulation, 2019. 12(6): p. 1381–1389.

17. Kasten, F.H., B. Maess, and C.S. Herrmann, Facilitated Event-Related Power Modulations during Transcranial Alternating Current Stimulation (tACS) Revealed by Concurrent tACS-MEG. eneuro, 2018. 5(3): p. ENEURO.0069-18.2018.

18. Kasten, F.H. and C.S. Herrmann, Transcranial Alternating Current Stimulation (tACS) Enhances Mental Rotation Performance during and after Stimulation. Frontiers in Human Neuroscience, 2017. 11.

19. Wischnewski, M. and D.J.L.G. Schutter, After-effects of transcranial alternating current stimulation on evoked delta and theta power. Clinical Neurophysiology, 2017. 128(11): p. 2227–2232.

20. Barone, J. and H.E. Rossiter, Understanding the Role of Sensorimotor Beta Oscillations. Frontiers in Systems Neuroscience, 2021. 15.

21. Engel, A.K. and P. Fries, Beta-band oscillations—signalling the status quo? Current Opinion in Neurobiology, 2010. 20(2): p. 156–165.

22. Kilavik, B.E., et al., The ups and downs of beta oscillations in sensorimotor cortex. Experimental Neurology, 2013. 245: p. 15–26.

23. Blanco Mora, D.A., et al., Toward methodologies for motor imagery enhancement: a tDCS-BCI study. Brain-Computer Interfaces, 2024. 11(3): p. 110–124.

24. Stančák, A., A. Riml, and G. Pfurtscheller, The effects of external load on movement-related changes of the sensorimotor EEG rhythms. Electroencephalography and Clinical Neurophysiology, 1997. 102(6): p. 495–504.

25. Tzagarakis, C., et al., Beta-band activity during motor planning reflects response uncertainty. The Journal of neuroscience : the official journal of the Society for Neuroscience, 2010. 30(34): p. 11270–11277.

26. Van Hoornweder, S., et al., Age and interlimb coordination complexity modulate oscillatory spectral dynamics and large-scale functional connectivity. Neuroscience, 2022.

27. Van Hoornweder, S., et al. Aging and Complexity Effects on Hemisphere-Dependent Movement-Related Beta Desynchronization during Bimanual Motor Planning and Execution. Brain Sciences, 2022. 12, DOI: 10.3390/brainsci12111444.

28. Alayrangues, J., et al., Error-related modulations of the sensorimotor post-movement and foreperiod beta-band activities arise from distinct neural substrates and do not reflect efferent signal processing. NeuroImage, 2019. 184: p. 10–24.

29. Bizovičar, N., et al., Decreased movement-related beta desynchronization and impaired post-movement beta rebound in amyotrophic lateral sclerosis. Clinical Neurophysiology, 2014. 125(8): p. 1689–1699.

30. Heinrichs-Graham, E., et al., Neuromagnetic evidence of abnormal movement-related beta desynchronization in Parkinson’s disease. Cereb Cortex, 2014. 24(10): p. 2669–78.

31. Salmelin, R., et al., Bilateral activation of the human somatomotor cortex by distal hand movements. Electroencephalography and Clinical Neurophysiology, 1995. 95(6): p. 444–452.

32. Stancák, A. and G. Pfurtscheller, Event-related desynchronisation of central beta-rhythms during brisk and slow self-paced finger movements of dominant and nondominant hand. Cognitive Brain Research, 1996. 4(3): p. 171–183.

33. Sisti, H.M., et al., Testing Multiple Coordination Constraints with a Novel Bimanual Visuomotor Task. PLOS ONE, 2011. 6(8): p. e23619.

34. Verstraelen, S., et al., Dissociating the causal role of left and right dorsal premotor cortices in planning and executing bimanual movements – A neuro-navigated rTMS study. Brain Stimulation, 2021. 14(2): p. 423–434.

35. Tashiro, S., et al., Probing EEG activity in the targeted cortex after focal transcranial electrical stimulation. Brain Stimul, 2020. 13(3): p. 815–818.

36. Rueda-Delgado, L.M., et al., Coordinative task difficulty and behavioural errors are associated with increased long-range beta band synchronization. Neuroimage, 2017. 146: p. 883–893.

37. R Core Team, R: A language and environment for statistical computing. 2021, R Foundation for Statistical Computing: Vienna, Austria.

38. RStudio Team, RStudio: Integrated Development for R. 2020, Rstudio: PBC, Boston, MA.

39. Smith, S.M. and T.E. Nichols, Threshold-free cluster enhancement: addressing problems of smoothing, threshold dependence and localisation in cluster inference. Neuroimage, 2009. 44(1): p. 83–98.

40. Mangiafico, S., rcompanion: Functions to support extension education program evaluation [R statistical package]. 2016.

41. Torrecillos, F., et al., Distinct Modulations in Sensorimotor Postmovement and Foreperiod β-Band Activities Related to Error Salience Processing and Sensorimotor Adaptation. The Journal of Neuroscience, 2015. 35(37): p. 12753–12765.

42. Rueda-Delgado, L.M., et al., Age-related differences in neural spectral power during motor learning. Neurobiol Aging, 2019. 77: p. 44–57.

43. Zaepffel, M., et al., Modulations of EEG Beta Power during Planning and Execution of Grasping Movements. PLOS ONE, 2013. 8(3): p. e60060.

44. Pastötter, B., F. Berchtold, and K.H. Bäuml, Oscillatory correlates of controlled speed-accuracy tradeoff in a response-conflict task. Hum Brain Mapp, 2012. 33(8): p. 1834–49.

45. Pistohl, T., et al., Decoding natural grasp types from human ECoG. Neuroimage, 2012. 59(1): p. 248–60.

46. Pozdniakov, I., et al., Online and offline effects of transcranial alternating current stimulation of the primary motor cortex. Scientific Reports, 2021. 11(1): p. 3854.

47. Gross, J., et al., Task-dependent oscillations during unimanual and bimanual movements in the human primary motor cortex and SMA studied with magnetoencephalography. Neuroimage, 2005. 26(1): p. 91–8.

48. Houweling, S., et al., Neural changes induced by learning a challenging perceptual-motor task. Neuroimage, 2008. 41(4): p. 1395–407.

49. Hu, K., et al., Effects of transcranial alternating current stimulation on motor performance and motor learning for healthy individuals: A systematic review and meta-analysis. Front Physiol, 2022. 13: p. 1064584.

50. Alekseichuk, I., M. Wischnewski, and A. Opitz, A minimum effective dose for (transcranial) alternating current stimulation. Brain Stimul, 2022. 15(5): p. 1221–1222.

51. Neri, F., et al., A novel tDCS sham approach based on model-driven controlled shunting. Brain Stimul, 2020. 13(2): p. 507–516.

52. Wischnewski, M., et al., Frontal Beta Transcranial Alternating Current Stimulation Improves Reversal Learning. Cerebral Cortex, 2020. 30(5): p. 3286–3295.

53. Oldfield, R.C., The assessment and analysis of handedness: The Edinburgh inventory. Neuropsychologia, 1971. 9(1): p. 97–113.

54. Oostenveld, R., et al., FieldTrip: Open Source Software for Advanced Analysis of MEG, EEG, and Invasive Electrophysiological Data. Computational Intelligence and Neuroscience, 2011. 2011(1): p. 156869.

