## Appendices for "The Causal Role of Beta Band Desynchronization: Individualized High-Definition Transcranial Alternating Current Stimulation Improves Bimanual Motor Control"

### A1. Hard- and software manufacturer information

| Hard- or software | Manufacturer information |
| --- | --- |
| Shaft encoded bimanual tracking task | A1230, Allegro microsystems, Manchester, United Kingdom |
| tACS stimulator: neuroConn DC-Stimulator Plus | Neurocare group AG, Munich, Germany |
| Ten20 paste | Weaver and Company, Aurora, United States |
| 64-channel BioSemi ActiveTwo EEG system | [Biosemi, Amsterdam, The Netherlands |
| MATLAB | The MathWorks Inc., Portola Valley, CA, USA |
| Signal Gel | Parker Laboratories, INC. Fairfield, United States |

A linear mixed model was constructed;

${Tracking Error Change}_{ijklmno}= \beta_{1}{Stimulation}_{ij}+ \beta_{2}{Block}_{ik}+ \beta_{3}\left( {Stimulation}_{ij}\times{Block}_{ik} \right)+ \beta_{4}{Laterality}_{il}+ \beta_{5}\left( {Stimulation}_{ij}\times{Laterality}_{il} \right)+\beta_{6}\left( {Block}_{ik}\times{Laterality}_{il} \right)+\beta_{7}\left( {Stimulation}_{ij}{\times Block}_{ij}\times{Laterality}_{il} \right)+\beta_{8} {Session}_{in}$	**Eq. 4**

Where $\beta_{1}$ to $\beta_{7}$ relate to the fixed effects of stimulation type, j, block, k, laterality, l, and their interactions, and the $\beta_{8}$ refers to the covariate for session, n. TE change values were aggregated to have one value per combination of stimulation, block, laterality, session and participant.

The final model, shown in **Table A4.1**., demonstrates significant effects of STIMULATION (F_2, 556.76_ = 11.305, p < 0.001), BLOCK (F_1, 544.35_ = 67.003, p < 0.001) and LATERALITY (F_2, 544.35_ = 4.213, p = 0.015), as well as for the covariate of SESSION (F_2, 556.80_ = 5.864, p = 0.003). No interaction effects remained after model building.

Table A4.1. Linear Mixed Effect Model for tACS effect on tracking error change.

|  | **Tracking Error Change** | | |
| --- | --- | --- | --- |
| *Predictors* | *Estimates* | *CI* | *p* |
| (Intercept) | -0.38 | -0.47 – -0.30 | **<0.001** |
| Stimulation [Individualized] | -0.15 | -0.21 – -0.09 | **<0.001** |
| Stimulation [Sham] | -0.06 | -0.12 – -0.00 | **0.040** |
| Block [Block 3] | -0.20 | -0.25 – -0.15 | **<0.001** |
| Laterality [Left] | 0.06 | 0.00 – 0.12 | **0.046** |
| Laterality [Right] | 0.08 | 0.03 – 0.14 | **0.005** |
| Session [s2] | 0.10 | 0.04 – 0.16 | **0.001** |
| Session [s3] | 0.08 | 0.02 – 0.14 | **0.013** |
| **Random Effects** | | | |
| σ^2^ | 0.09 | | |
| τ_00_ _ID_ | 0.02 | | |
| ICC | 0.22 | | |
| N _ID_ | 36 | | |
| Observations | 588 | | |
| Marginal R^2^ / Conditional R^2^ | 0.129 / 0.322 | | |

Tukey-corrected post-hoc tests for the effect of STIMULATION show that individualized tACS resulted in a greater improvement on the BTT than sham tACS (estimate = -0.09, t_555_ = -2.848, p = 0.013) and fixed tACS (estimate = -0.15, t_555_ = -4.727, p < 0.001). Concerning block (Table 2), as to be expected, BTT improvement was higher in block 3 than block 2. Post-hoc tests of the LATERALITY effect demonstrated that there was a significant difference between BTT conditions wherein both hands had to move at equal speeds, compared to conditions where the right hand had to move faster, with the latter conditions showing the greatest improvements with respect to block 1 (estimate = -0.08, t_545_ = -2.823, p = 0.013).

**Figure A4.1.** visually shows the main effects. As can be seen in this figure, there are some outliers with the most apparent ones being present in the individualized tACS condition. When rerunning the aforementioned model without outliers, defined by a distance larger than 3 standard deviations of the mean, the effect of STIMULATION and the associated post-hoc tests remained identical in terms of significance and interpretation.

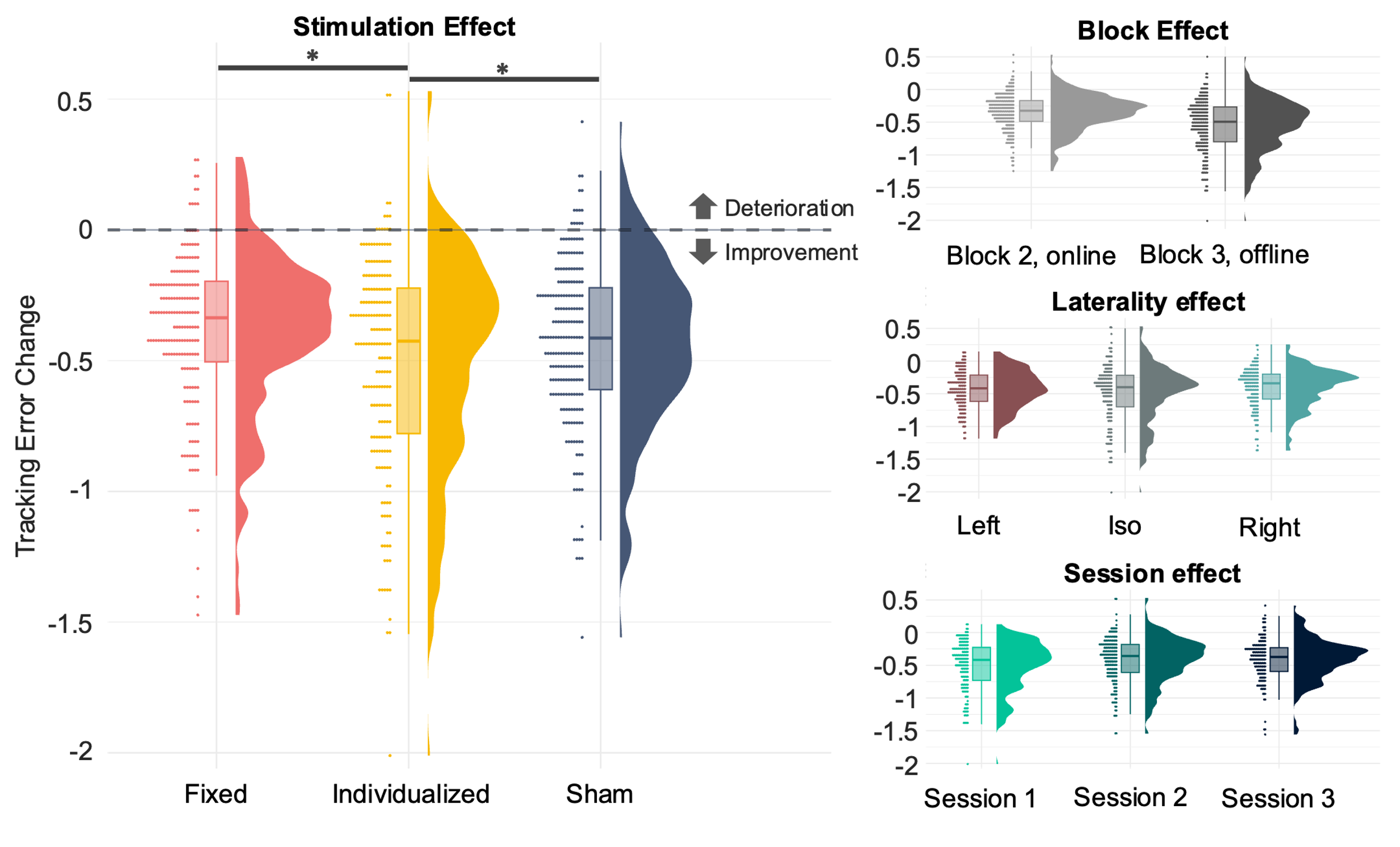

**Figure A4.1.** The effect of stimulation, block, laterality and session on mean tracking error change. The left panel shows the effect of stimulation, with greater increases in mean tracking error as a result of individualized tACS. The right upper panel shows the effect of block, with a larger tracking error improvement in block 3, compared to block 1.

### Appendix 5. Full Threshold Free Clustering Enhanced (TFCE) time course of online stimulation effect

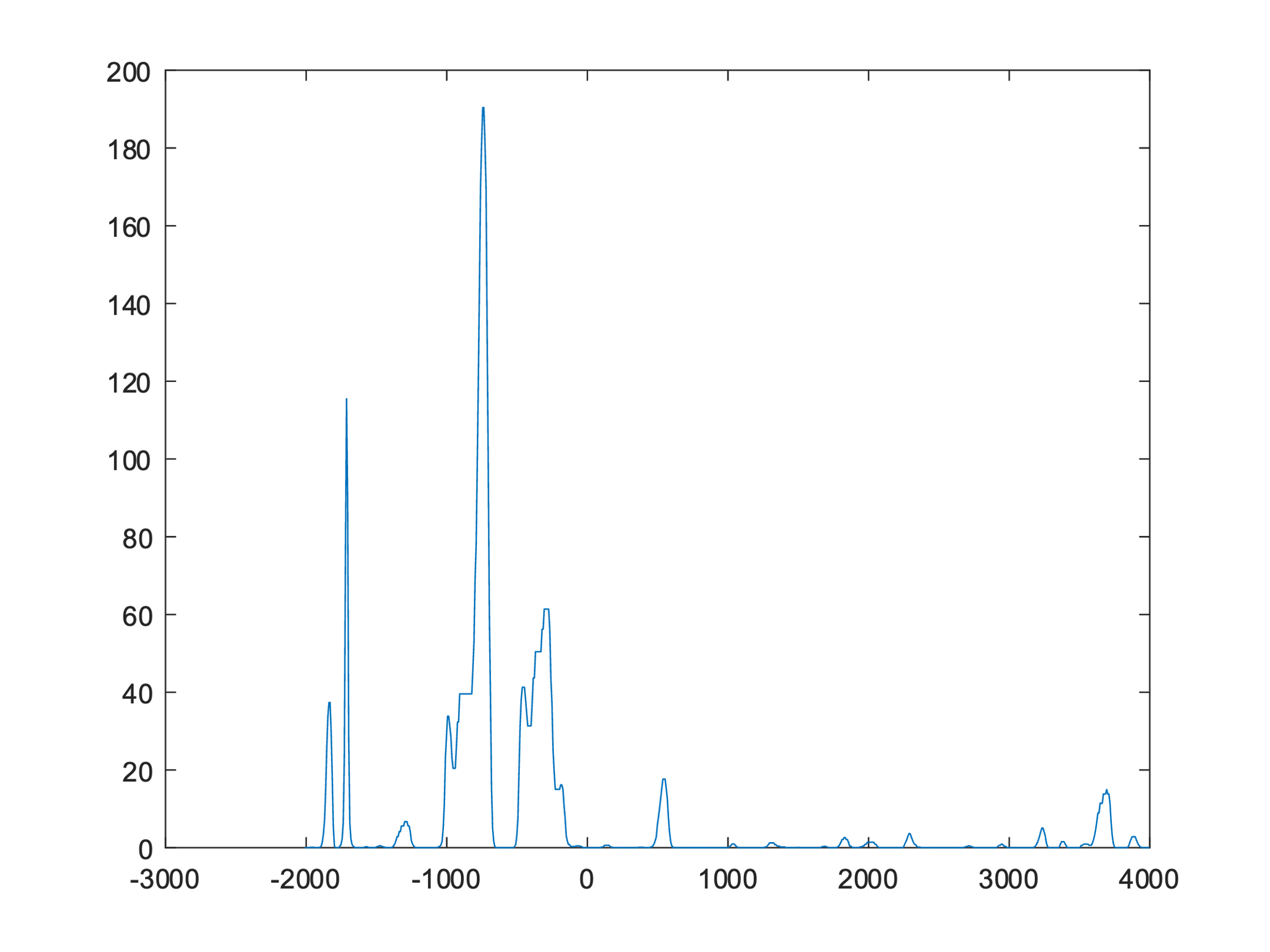

**Figure A5.** TFCE time course of the online stimulation effect on MRβD (fixed vs. individualized tACS). The Y axis denotes TFCE values, the X-axis denotes time in ms. As can be seen, the largest TFCE values occur during the planning phase (-2000 to 0 ms), whereas values TFCE during motor execution are low.

### A6. Comparison of the right and left EEG sensors to predict bimanual motor control improvements

Previous work suggests that the right sensorimotor cortex is particularly sensitive to increased motor task demands. To further test this with the available data, we aimed to see if perhaps data from EEG sensor C3 (left hemisphere) was better able to predict tracking error change than EEG sensor C4.

We ran the model defined in **Section 3.4.** using EEG data from sensors C3 or C4 and compared both models. The model used in the manuscript, incorporating sensor C4, best explained changes in tracking error, as shown in **Table A6**.

**Table A6.** Comparison of TE Change prediction using peak planning MRβD extracted from the right (C4) and left (C3) sensorimotor regions.

| Model | R^2^_marginal_ | R^2^_conditional_ | AIC | BIC |
| --- | --- | --- | --- | --- |
| TE Change = ∆(peak MRβD **C4**) + Stimulation | 0.201 | 0.300 | 36.02 | 51.28 |
| TE Change = ∆(peak MRβD **C3**) + Stimulation | 0.137 | 0.252 | 43.89 | 59.15 |
